# Modular, cement-free, customized headpost and connector-chamber implants for macaques

**DOI:** 10.1101/2022.11.09.515849

**Authors:** Eleni Psarou, Julien Vezoli, Marieke L. Schölvinck, Pierre-Antoine Ferracci, Yufeng Zhang, Iris Grothe, Rasmus Roese, Pascal Fries

## Abstract

**Background:** Neurophysiological studies with awake macaques typically require chronic cranial implants. Headpost and connector-chamber implants are used to allow head stabilization and to house connectors of chronically implanted electrodes, respectively.

**New Method:** We present long-lasting, modular, cement-free headpost implants made of titanium that consist of two pieces: a baseplate and a top part. The baseplate is implanted first, covered by muscle and skin and allowed to heal and osseointegrate for several weeks to months. The percutaneous part is added in a second, brief surgery. Using a punch tool, a perfectly round skin cut is achieved providing a tight fit around the implant without any sutures. We describe the design, planning and production of manually bent and CNC-milled baseplates. We also developed a remote headposting technique that increases handling safety. Finally, we present a modular, footless connector chamber that is implanted in a similar two-step approach and achieves a minimized footprint on the skull.

**Results:** Twelve adult male macaques were successfully implanted with a headpost and one with the connector chamber. To date, we report no implant failure, great headpost stability and implant condition, in four cases even more than 9 years post-implantation.

**Comparison with Existing Methods:** The methods presented here build on several related previous methods and provide additional refinements to further increase implant longevity and handling safety.

**Conclusions:** Optimized implants can remain stable and healthy for at least 9 years and thereby exceed the typical experiment durations. This minimizes implant-related complications and corrective surgeries and thereby significantly improves animal welfare.

**Highlights:**

- Long-lasting titanium implants for non-human primates
- Refined implantation techniques that reduce post-operative complications
- Minimized, footless connector chamber to house connectors of chronic arrays
- Twelve adult male macaques were implanted with long-lasting headpost implants
- Headpost implants so far without failure and with longevity up to > 9 years

## 1 Introduction

Understanding the primate brain requires neurophysiological studies with awake behaving macaque monkeys. The majority of these studies so far requires head fixation that greatly eases the precise monitoring of eye position, and that is required for most recording approaches. The ability of head fixation also allows the experimenter to safely provide wound care to awake animals.

While conducting animal research, it is an ethical imperative to comply with the 3R principles: replacement, reduction, refinement. Refinement of procedures in the field of awake macaque monkey research is particularly challenging, because most studies in the field typically include a very low number of animals. Therefore, even small refinements obtained in one laboratory should be shared and disseminated. This could help many researchers refine their techniques and promote the welfare of many experimental monkeys.

Research on the neural substrate of many higher cognitive functions builds on the ability of macaque monkeys to perform complex cognitive tasks. Such tasks often require extended training periods that can last up to several months. Moreover, in the case of studies with awake macaques, refinement can also lead to reduction. Given their long lifespan, one animal can often participate in several subsequent projects as long as the animal and the implants are in good health. Therefore, there is great need for long-lasting implants that can stay healthy over extended time periods.

In the last two decades, there has been a considerable effort to refine headpost implants and their implantation techniques. Several improvements have led to higher success rates and longer implant lives, e.g. through the customization of implant shapes (Adams et al., 2007; Chen et al., 2017; Overton et al., 2017), the use of more biocompatible materials (Adams et al., 2007; Lanz et al., 2013; Overton et al., 2017), the use of coatings to enhance osseointegration (Chen et al., 2017; Lanz et al., 2013), or the use of two-step implantation approaches (Betelak et al., 2001; Blonde et al., 2018). Here, we present our approach that builds on previously reported methods (Adams et al., 2007; Johnston et al., 2016; Lanz et al., 2013; Overton et al., 2017), and we describe additional refinements to the entire head fixation technique, including the implant itself, the surgical procedures and the everyday handling. We also present a connector-chamber implant that houses chronic electrode connectors. This connector chamber is inspired by our headpost approach and aims to reduce the overall footprint of the implant on the skull, and to facilitate its osseointegration.

Briefly, 1) we developed long-lasting modular implants that are implanted in a refined two-step approach, 2) we provide detailed protocols of our implant design, planning and production procedures, and of our implantation techniques, 3) we share the 3D models of the implants and tools we developed, so any lab can reproduce them, and 4) we present the results from twelve adult male macaque monkeys that were implanted with these techniques. Importantly, we experienced no implant failure and found the implants to last up to more than 9 years, i.e. until today.

## 2 Materials and methods

### 2.1 Animals

Twelve male monkeys (*Macaca mulatta*) were implanted with headpost implants (see details in **Table 1**) and one with the connector chamber (Monkey C). All procedures and housing conditions complied with the German and European law for the protection of animals (EU Directive 2010/63/EU for animal experiments). All surgical and experimental methods were approved by the regional authority (Regierungspräsidium Darmstadt) under the following permit numbers: F149/01, F149/07, F149/08, F149/1003, F149/1007, F149/1008, F149/1010, and F149/2000.

**Table 1.** Monkey Information.

| Monkey | Age* (year) | Weight* (kg) |
| --- | --- | --- |
| Monkey L | 8 | 11.0 |
| Monkey Sk | 8.8 | 11.5 |
| Monkey D | 8.7 | 8.3 |
| Monkey G | 8 | 11.2 |
| Monkey Hu | 7 | 11.5 |
| Monkey Ch | 12.3 | 16.0 |
| Monkey T | 7.9 | 13.1 |
| Monkey St | 12.5 | 15.0 |
| Monkey M | 6.9 | 11.9 |
| Monkey H | 15.4 | 15.0 |
| Monkey C | 15.3 | 14.0 |
| Monkey K | 15.4 | 13.0 |

\* At the time of the headpost baseplate implantation. Animals are listed in the order of implant longevity until today or until sacrificed or deceased, as listed in Table 7.

### 2.2 A two-piece headpost

We developed a cement-free, two-piece headpost that consists of a baseplate and a top part that are implanted in two separate surgeries. **Figure 1** presents a graphical overview of our headpost methods. The baseplate is implanted first (**Fig. 1A)**. It is customized to follow the skull surface of the individual monkey and is anchored onto the bone exclusively by means of titanium bone screws. The screw length is adjusted to match the skull thickness. At the end of this surgery, the baseplate is covered with muscle and skin, and the surgical site is allowed to heal for several weeks. During this period, the sterile conditions established by the closing of the skin provides optimal conditions for osseointegration. Following an adequate waiting period (see **section 2.3.2**), the percutaneous part of the implant is added (**Fig. 1B**). In a short surgery, the central plate of the baseplate is exposed, and the top part is secured onto it using a screw (top-part screw). Finally, we also developed a headpost holder that allows remote headposting for increased safety during monkey handling (**Fig. 1C)**.

**Figure 1.**
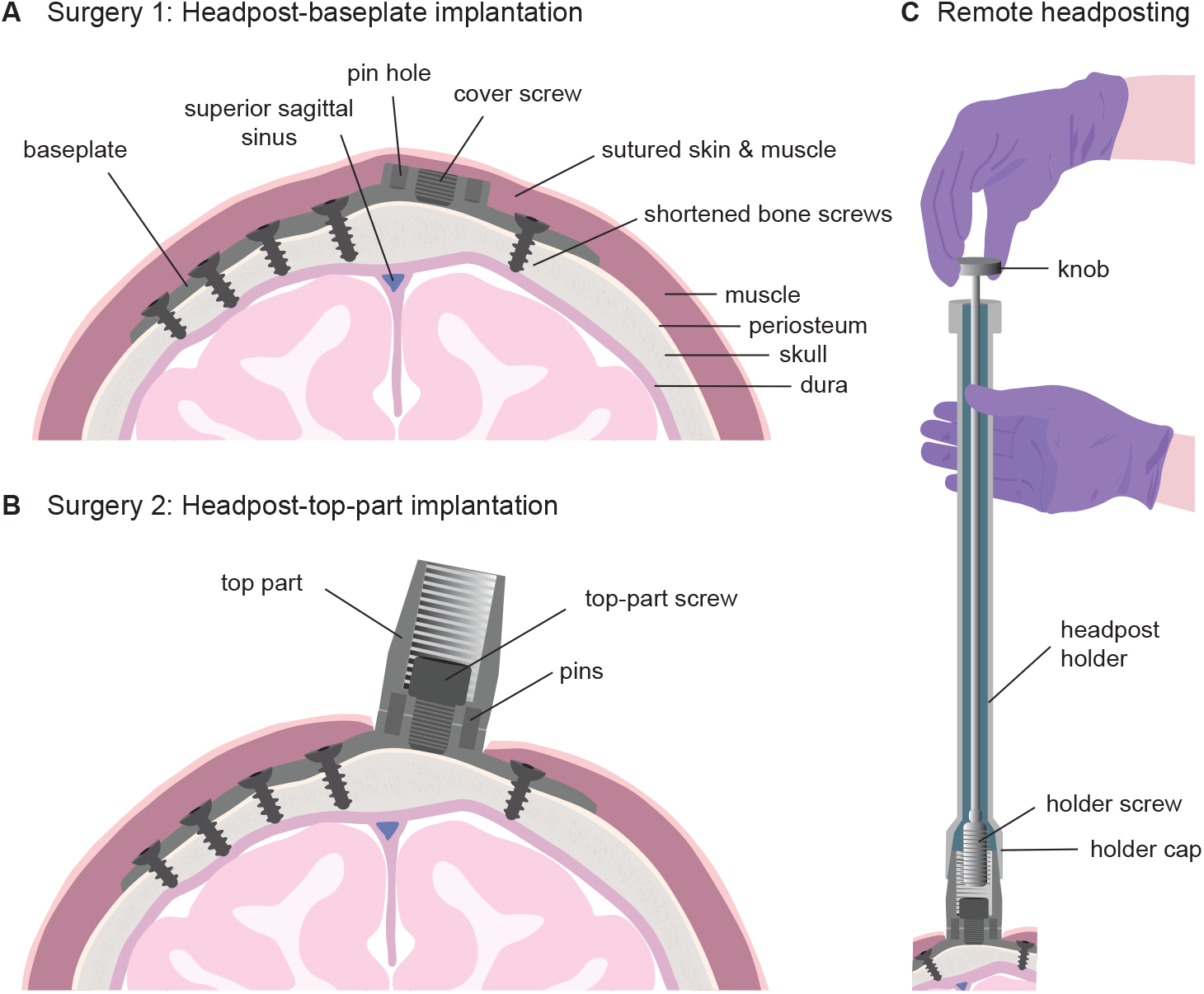
Illustration of headpost methods. **A**) The baseplate is implanted first, secured to the skull with titanium bone screws, whose length has been adjusted to match the underlying bone thickness. At the end of this surgery, the baseplate is covered with muscle and skin, and the surgical site is allowed to heal for several weeks. **B**) The top part is added in a separate short surgery. A circular cut is performed and the top part is secured onto the baseplate with a central screw. Two pins prevent the top part from rotating. **C**) The headpost holder allows the experimenter to remotely fixate the animal’s head. By turning a knob at the proximal end of the headpost holder, the holder screw at the distal end enters into the thread of the top part and thereby fixes the holder to the implant.

#### 2.2.1 Headpost baseplate

Two baseplate versions were used that differed in how they were shaped to follow the skull surface: one version was computer-numerical-control (CNC) milled in a flat shape and then customized by manual bending (referred to as “bent version”, **Fig. 2A**); another version was CNC-milled directly to follow the skull shape (referred to as “milled version”, **Fig. 2B-C**). Both versions are based on an overall similar design. Nine monkeys have been implanted with the bent version and three with the milled one (**Table 2**).

**Figure 2.**
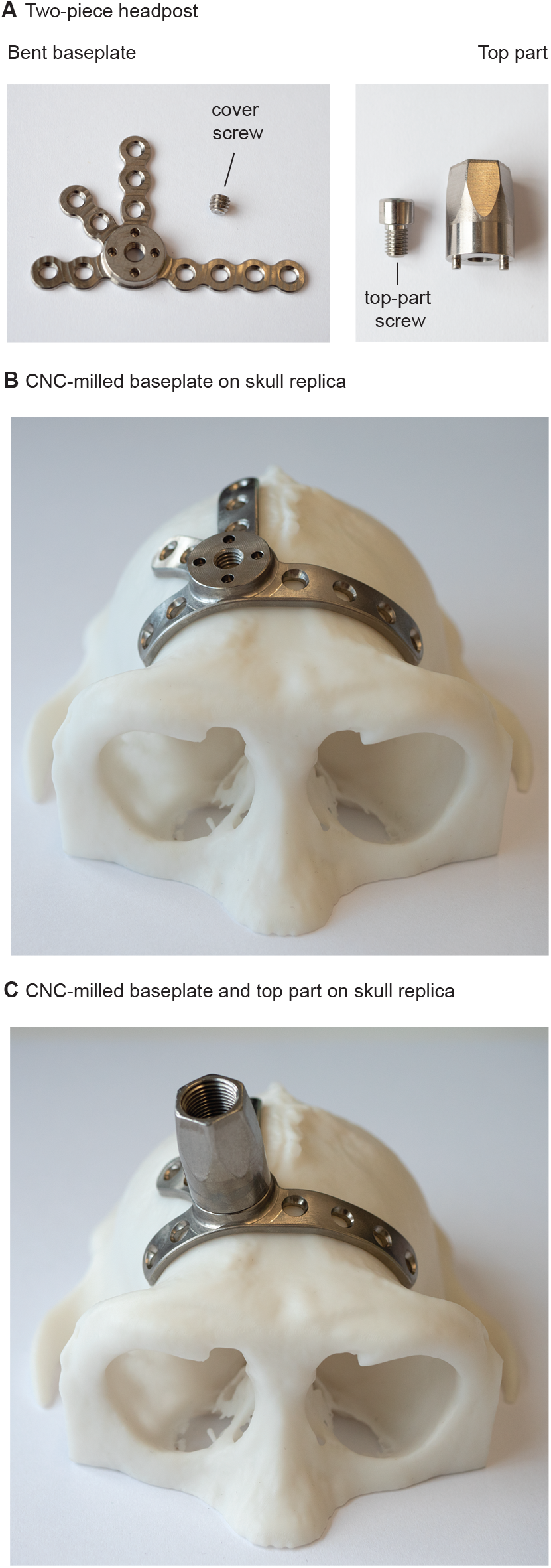
The modular headpost implant. **A**) The main parts of the headpost are shown: on the left, a baseplate (flat precursor of bent version shown as example) with the cover screw, and on the right, a top part and the titanium screw that secures the two parts together (top-part screw). **B**) The CNC- milled version of the baseplate (frontal version) is shown on top of the 3D printed skull replica of Monkey H. **C**) The top part is added on the baseplate.

**Table 2.** Software used for the planning of the headpost baseplate.

| Monkey | Software | Baseplate Version |
| --- | --- | --- |
| Monkey L | Brain Voyager QX | bent, frontal |
| Monkey Sk | BrainVoyager QX | bent, frontal |
| Monkey D | BrainVoyager QX | bent, frontal |
| Monkey G | Brain Voyager QX | bent, frontal |
| Monkey Hu | Brain Voyager QX, Image J | bent, frontal |
| Monkey Ch | Brain Voyager QX, ImageJ | bent, frontal |
| Monkey T | Brain Voyager QX | bent, frontal |
| Monkey St | 3D Slicer 4.10.0 | bent, frontal |
| Monkey M | Brain Voyager QX, ImageJ | bent, frontal |
| Monkey H | 3D Slicer 4.10.0, Geomagic Design X, SOLIDWORKS | milled, frontal |
| Monkey C | 3D Slicer 4.10.0, Geomagic Design X, SOLIDWORKS | milled, central |
| Monkey K | 3D Slicer 4.10.0, Geomagic Design X, SOLIDWORKS | milled, central |

Both versions were produced from titanium Grade 2. The advantage of the bent version is that it is easy, cheap and fast to produce in larger numbers, and can be shaped to the individual skull of a monkey whenever its implantation is planned. The flat precursor of the bent baseplate (**Fig. 2A**) can be produced on a 3-axis CNC machine. The advantage of the milled version is that it provides a near perfect fit to the skull (**Fig. 2B-C**), which cannot be achieved by manual bending. The milled baseplate requires a 5-axis CNC-milling machine.

The baseplate contains a central plate, onto which the headpost top part is later mounted, and several “legs” that extend radially from this plate. The overall shape of the baseplate with its legs varied depending on the brain areas of interest. As described in **Table 2**, ten monkeys were implanted with a baseplate in the most anterior part of the skull (frontal version), and two with a more centrally located implant (central version).

The frontal baseplate version **(Fig. 3A)** was designed to allow later access to almost the entire left hemisphere (except the most frontal areas), and to occipital, temporal and part of parietal areas on the right hemisphere. The central version **(Fig. 3B)** was designed to allow later access to both frontal and occipital areas of the left hemisphere, and to occipital areas of the right hemisphere.

**Figure 3.**
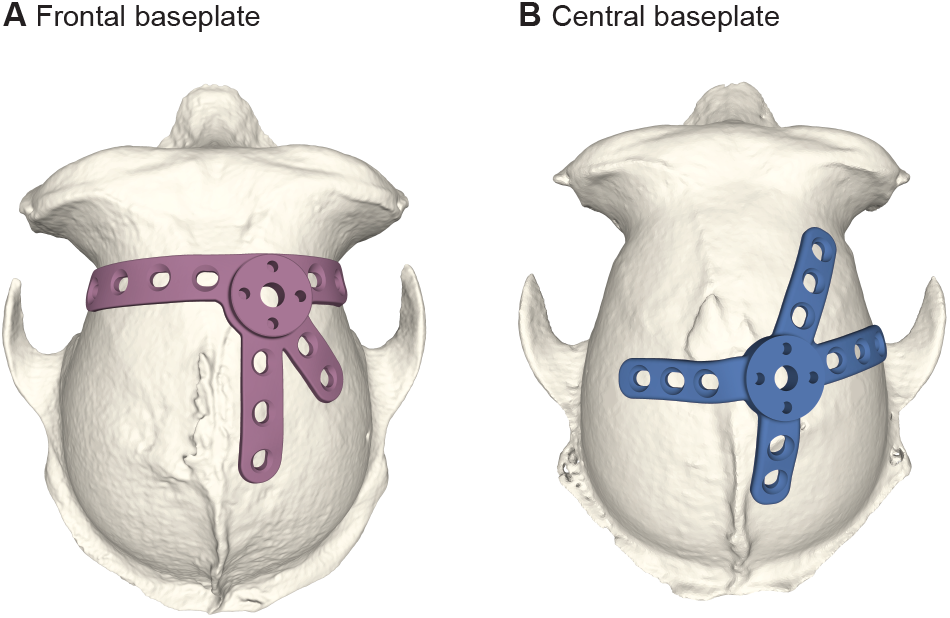
Versions of the CNC-milled headpost baseplate. **A**) A frontal baseplate (Monkey H) and, **B**) a central baseplate (Monkey C) allow access to different brain areas.

In planning the baseplate position and implantation, one needs to consider the underlying anatomy. The most anterior legs of the frontal baseplate version ran parallel to the supraorbital ridge, and we chose to have them run several millimeters behind this ridge. This had several reasons: 1) Screws placed even further anterior can connect to the frontal sinus, which can be a source of infection (though we note that several other labs have successfully implanted similar designs on top of the frontal ridge: Lanz et al. (2013), Overton et al. (2017), Adams et al. (2007), Adams et al. (2011), Ortiz-Rios et al. (2018)); 2) This position avoids the screws entering the eye socket; 3) This position coincides with the coronal section through the skull with a particularly small radius, whereby the baseplate legs can “grab” particularly effectively around the skull, which in turn provides optimal anchoring in the bone at a relatively large angle relative to the pulling force on the headpost (note the two most lateral bone screws in **Fig. 4C-D**). The leg extending in the anterior-posterior direction, along the midline, was always placed slightly away from the midline, in order to avoid the screws damaging the superior sagittal sinus.

**Figure 4.**
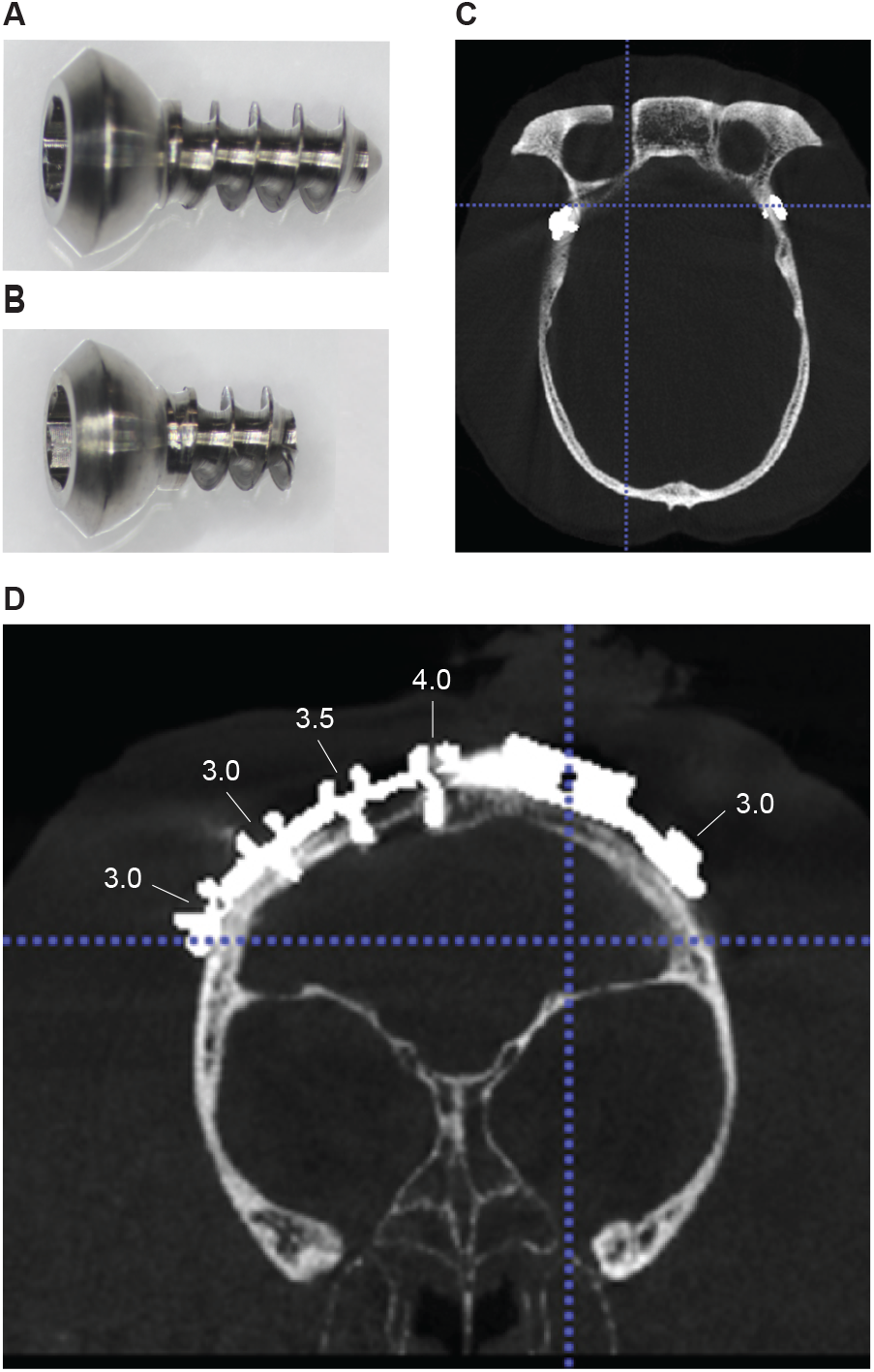
Length-adjusted bone screws. The original self-tapping, titanium bone screws (**A**) were cut (**B**) in order to match the thickness of the underlying bone. **C-D**) Post-operative CT scan of Monkey St that shows the successful choice of screw lengths at the anterior legs of the baseplate implant. The thread length of the adjusted screws is indicated. Panel C shows the thread of the two more lateral screws of which the head is obvious in D.

An earlier version of the frontal baseplate was anchored to the skull with twelve bone screws (Monkey Sk, Monkey D and Monkey T). Later, one screw hole was removed from the leg running between the most anterior and the one parallel to the midline. This allowed access to more brain areas of the right hemisphere. Nine monkeys received a baseplate with eleven bone screws.

#### 2.2.2 Headpost top part

The headpost top part (**Fig. 2A** and **2C**) is a separate CNC-milled piece (requiring 3 or more axes), made of titanium Grade 5, which is mounted on the baseplate (made from titanium Grade 2) in a second surgery. It is the percutaneous part of the implant that can be secured by the headpost holder to allow head fixation. The top-part is fixed to the central plate of the baseplate with a commercially available screw of size M5, whose head diameter is reduced in-house to fit within the top part. We refer to it as ‘top-part screw’ and it is also made of titanium Grade 5. Two stainless steel pins on opposite sides of the base of the top part fit into respective pinholes on the central plate of the baseplate **(Fig. 2A)**. Those pins allow to define the precise orientation of the top part before its implantation, and they prevent any rotation of the top part after implantation. Due to a problem that occurred in one animal (see Results, **section 3**), we added two additional, spare, pin holes on the baseplate.

As can be seen in **Figure 2C**, the top part contains on its inside a 9 mm diameter hole with a screw thread, and on its outside a hexagonal surface that narrows conically towards the top. These features allow a connection to the headpost holder, as described below under **section 2.4**.

#### 2.2.3 Skull reconstruction

The implant planning starts with the acquisition of a preoperative computerized tomography (CT) scan in order to create a precise model of the individual animal’s skull. The 3D model is then printed and used to guide individualized implant planning and production. Even though a skull model can be extracted from MRI scans, CT scans provide more precise information about the bone structure, yielding to an easier and more detailed skull reconstruction.

The scan is performed under ketamine-medetomidine anesthesia and in case the animal is placed in a stereotaxic frame, the anesthesia is combined with NSAIDs and the application of lidocaine ointment on the tip of the ear-bars. Seven monkeys were scanned in a Brilliance 6 scanner (Philips, Amsterdam, Netherlands) and five in a ProMax 3D Mid scanner (Planmeca Oy, Helsinki, Finland). For a detailed summary of the scanning parameters used per monkey refer to **Suppl. Table 1**.

The ProMax is a CBCT (Cone Beam Computed Tomography) scanner typically used in dentist and ear-nose-throat applications, where it provides an extended view of the maxillo-facial region. We used a slightly adapted version of such a system to obtain CTs of nearly the entire macaque skull in one volume. The system has a similar size as a surgical microscope, and is similarly mounted on a mobile platform and can thereby be easily used in a research setting (though the room needs to be specially equipped for the use of X-ray radiation). After positioning of the animal, the CT measurement is obtained within less than one minute; the calculation of the CT reconstruction on a regular PC takes few minutes.

To make optimal use of the high resolution of the CT scans, we recommend to place the animal’s head into a stereotaxic frame. Also, such a frame provides ear bars and eye bars. The presence of ear bars in the CT images can be helpful in the determination of the inter-aural line (an imaginary line connecting the tips of the ear bars). The eye bars are used to define the inferior-orbital ridge. The inter-aural line together with the inferior-orbital ridges can be used to define the orbitomeatal plane, also referred to as Frankfurt baseline plane (Dubowitz and Scadeng, 2011) or Frankfurt zero plane. In the Brilliance 6 scanner, the head can be positioned in an MRI compatible stereotaxic frame that allows artifact-free CT scans.

In the ProMax (Planmeca) scanner, we have typically used a custom-built stereotaxic frame. This stereotaxic frame positioned the head above the lateral bars, which allowed the CT scan to contain most of the skull. The resulting volume did not reach down to the skull base, but it did contain all skull features relevant for the Frankfurt-zero alignment, namely the ear canals, the occipital ridge, and the complete eye sockets including the lower margin of the eye socket (see example scan in **Fig. 4C-D**). We strongly recommend to place a CT-visible marker on one side of the animal’s head to avoid ambiguities with regard to the orientation of the images (see MRI-marker in **Fig. 5A-C**). We used one of the CT-visible markers that came with the MRI-compatible stereotaxic frame (see next paragraph).

**Figure 5.**
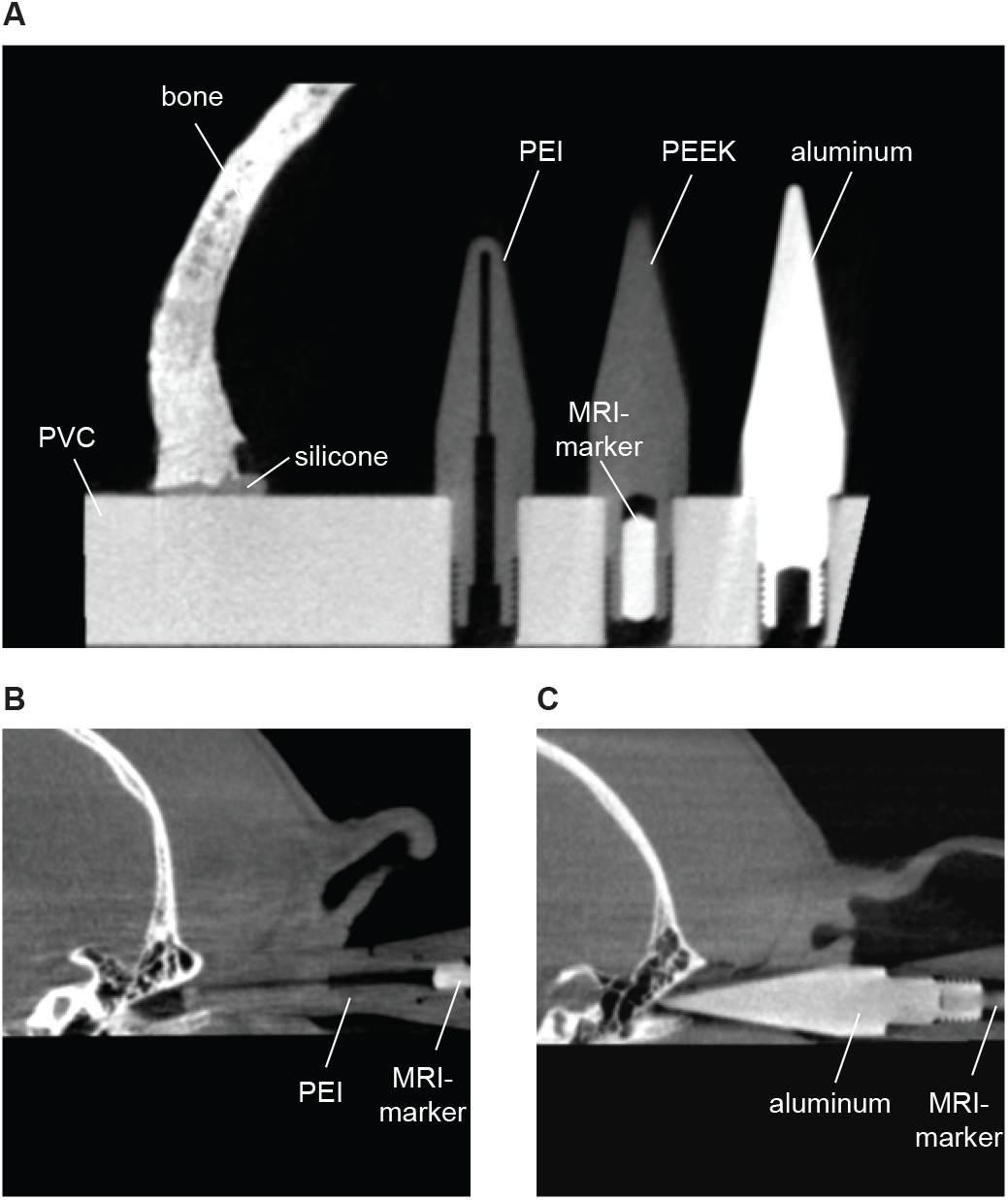
Comparison of X-ray contrast of different materials. **A**) Ear-bar tips made from different materials and skull bone are shown for comparison. Aluminum gives stronger X-ray contrast compared to polyetherimide (PEI) and polyether ether ketone (PEEK). **B**) CT scan of Monkey St with the PEI ear bar. **C**) CT scan of Monkey C with the aluminum ear-bar tip mounted on the handle of the MRI- compatible PEI ear bar. The MRI marker was placed on one side only, to avoid ambiguities in the left- right orientation of the CT images.

As mentioned above, the ear bars of the stereotaxic frame can be used in determining the interaural line in the CT scans. The commercially available ear-bars of our MRI-compatible stereotaxic frame (Model 1430M MRI Stereotaxic Instrument, David Kopf instruments, Los Angeles, California, USA) were made of polyetherimide (PEI) which does not produce artifacts. However, the PEI also does not provide very good X-ray contrast (**Fig. 5A-B**), and the tips of the ear bars were relatively blunt/round (**Fig. 5A-B**), and not identical to the ear bars used later during implantation surgeries. Therefore, we used the opportunity to exchange the tips on those ear bars. We exchanged the tips with custom-built tips made of aluminum. The aluminum gives excellent X-ray contrast (**Fig. 5A** and **5C**), and at the same time minimizes stray artifacts, and it was easy to produce tip diameters optimized for monkey ear canals and identical to those used during implantation surgeries. The use of identical tip diameters during CT scanning and surgery improves the alignment between the two head fixations.

From the CT based 3D volume, the skull was segmented using either Brain Voyager QX (Goebel et al., 2006) or 3D Slicer (Fedorov et al., 2012). **Table 2** summarizes the software packages used per monkey for implant planning.

#### 2.2.4 Baseplate shaping to the skull

##### 2.2.4.1 Shaping of the baseplate with manual bending

A 3D printed replica of the monkey’s skull is used as template in order to prepare the manually bent baseplate preoperatively (for a similar approach see Overton et al. (2017)).

In the case of the anterior baseplate version, simple landmarks on the skull (like the supraorbital ridge and midline) can be used to guide the positioning of the implant. We used a simple paper version of the baseplate. We placed this paper onto the 3D skull replica and gently pushed it down, so it adapted to the shape of the skull. We then shifted it around to find the optimal position. In doing so, we took into account the considerations mentioned above, under **section 2.2.1**, namely 1) that the baseplate optimally “grabs” around the part of the skull with a relatively small radius, and 2) that the screws avoid the supraorbital ridge (and thereby the underlying sinus), the eye sockets, and the superior sagittal sinus in the midline. Note that we did not attempt the central plate to be positioned precisely on the midline, which would allow a vertical top part (see **section 4.7.1**).

Once the final location had been decided, an outline of the baseplate was drawn on the skull replica that later on guided the manual bending of the titanium implant. Careful bending of the baseplate legs to achieve an optimal fit can take up to several hours. For this reason, this procedure should be completed before the surgery. It is also much easier and safer to handle and bend the implant in a non-sterile setting. For the manual bending, we used tools from DePuy Synthes (Raynham, MA, U.S.A; see **Suppl. Table 2**).

The bending starts with the part of the legs proximal to the central plate. Once the proximal parts of each of the legs have been bent to fit to the skull as good as possible, bending proceeds to more distal parts of the legs. Each step of the bending procedure should progressively approximate the optimal shape in small steps, rather than using multiple forward-backward bends, which can lead to material weakening and breakage. In doing so, the baseplate will need to be repeatedly placed against the skull replica to visually check its fit.

We found that the central plate with the most proximal parts of the legs constitute a relatively large part that cannot be bent and sufficiently adapted to the skull, such that gaps of 1-2 mm in some animals could not be avoided. This and an improvement of the overall fit to the skull were the main motivations to move to the milled baseplate, described in the next paragraph.

##### 2.2.4.2 Shaping of the baseplate with CNC-milling

Three monkeys were implanted with baseplate implants that were shaped to follow the individual skull geometry using CNC-milling (**Table 2**). Similar to the bent version described above, also the milled baseplates feature 1) four radially extending legs with eleven screw holes and, 2) a central plate that allows the top part to be mounted on it (the top part is identical to the one used with the manually bent baseplate).

Monkey H was implanted with the frontal version of the CNC-milled baseplate (**Fig. 3A**). While planning this implant, we aimed to produce a baseplate with similar design to the bent frontal version. To this end, our first implant planning steps were identical to the planning of the bent version (see **section 2.2.4.1**): the implant location was chosen using the skull replica and a paper version of the baseplate. The outline of the baseplate and each screw hole was then drawn onto the skull model. At this stage, one needs to translate the implant location into coordinates on the skull segmentation that can subsequently guide the implant design in the planning software. To do so, we drilled small holes into the 3D skull replica at the center of each screw hole and then acquired a CT volume of it. In the resulting volume, one could easily see the screw holes. This volume was then aligned to Frankfurt- zero and segmented. By overlying this model with the segmentation obtained from the original CT-volume of the monkey’s head, we could infer the target location of each screw and thus, the overall location and shape of the implant legs. Note that Ahmed et al. (2022) present a way to perform virtual bending of a headpost implant that could simplify this procedure.

Monkey C and Monkey K were implanted with a central CNC-milled baseplate version (**Fig. 3B**) whose position and shape were planned in a different way. First, the coordinates of the brain areas of experimental interest were estimated in the skull segmentation, and then the baseplate legs were designed to allow later access to those areas.

In all cases, the skull was segmented from the CT volume using 3D Slicer (Fedorov et al., 2012). The resulting skull model was imported into the Geomagic Design X software (https://www.oqton.com/geomagic-designx/) for further processing that facilitated the overall planning procedure. An area of interest on the skull was chosen and isolated (Lasso selection). Then, the isolated surface was fitted with a mesh using the function “Mesh Fit”.

An alternative to the Geomagic Design X software might be the FreeCAD software, an open-source parametric modeler (https://www.freecad.org/). It has not been used in the current experiments, but offers similar functionality. FreeCAD can be used for repairing and smoothing mesh data (with the “Mesh Workbench”), fitting a parametric surface to the skull model (with the “Surface Workbench”), and creating parametric models of solid parts (with the “Part Design Workbench”). Note that FreeCAD can also be an alternative for the SOLIDWORKS software.

In the case of Monkey C and Monkey K, two surfaces were created: a) a detailed one that closely follows the geometry of the skull (resolution by allowable deviation: 0.1 mm) and, b) a smoother version of it (allowable deviation: 1 mm).These two reconstructed surfaces were exported as parasolid surfaces which were then imported into SOLIDWORKS and used for the next planning steps. The detailed one was used to create the lower surface of the baseplate which provided a near perfect fit of the implant to the skull. The upper surface of the implant legs was based on the smoothed surface plus an offset to realize the thickness of the legs. This procedure led to uneven thickness along the implant legs (**Table 3**; Monkey C, Monkey K). To avoid weak points on the implant, the thickness of very thin parts was increased so that it was always more than 1.26 mm.

**Table 3.** Thickness of the different headpost baseplate versions.

| Baseplate version | Implant thickness (mm) |  |  |
| --- | --- | --- | --- |
|  | Leg |  | Central plate |
|  | *min.* | *max.* |  |
| bent | 1.70 | 1.70 | 4.5 |
| milled, Monkey H | 1.70 | 1.75 | 3.6 |
| milled, Monkey C | 1.67 | 2.80 | 4.0 |
| milled, Monkey K | 1.26 | 2.50 | 4.0 |

In the case of Monkey H, only the detailed surface was used for the implant planning. An offset was applied on this surface in order to achieve the desired implant thickness, and then, its upper surface was smoothed to avoid sharp features that could irritate the overlying muscle or skin.

##### 2.2.4.3 CNC-milling of the headpost baseplate

The CNC-milled headpost baseplates were in-house produced using a 5-axis CNC machine. During this process, material is progressively removed from a block of titanium until the final result is achieved. A crucial step is the clamping and thus, proper fixation of the titanium block throughout the milling procedure. We devised two different clamping approaches illustrated in **Suppl. Fig. 1** and **Suppl. Fig. 2**, respectively. We recommend the approach illustrated in **Suppl. Fig. 2**, because it avoids the need to plan and mill an extra piece, and it avoids potential imprecisions incurred by re-clamping (see **Suppl. Fig. 1** legend for details).

#### 2.2.5 Planning and preparation of bone screws

The baseplate is secured to the skull only by means of titanium screws. Eleven monkeys were implanted with commercially available screws (Crist Instrument Company, Inc., Hagerstown, Maryland, USA) and one (Monkey Ch) with in-house made titanium bone screws.

The commercially available bone screws (**Table 4**) came with a total length of 8.1 mm, a thread length of 5.5 mm and a thread diameter of 2.6 mm. In our experience, the bone at many parts of the macaque skull can be thinner than 5.5 mm. Therefore, in nine monkeys, we used the procedure described in the following.

**Table 4.** List of implant parts and implantation instruments used in the headpost-baseplate implantation.

| Implanted parts | Material | Production/ source |
| --- | --- | --- |
| baseplate | titanium Grade 2 | in-house |
| self-tapping bone screws | titanium | Crist Instrument |
| cover screw | stainless steel, A2 | commercially available |
| Implantation instruments | Specifications |  |
| manual drill system: |  |  |
| manual drill & drill guide with stop | Ø1.8 mm drill bit |  |
| electric drill (in case of bone smoothing) | electric pen drive,<br>Ø3-4 mm drill bit | DePuy Synthes |

We determined the optimal length of each screw preoperatively. This allowed to pre-adjust a drill stop for each screw (see below), and it removed the need to manually measure the bone thickness during the surgery. This screw-length adjustment was done on a lathe. **Figure 4 (A-B)** shows an original and an example shortened screw. We prepared screws with several thread lengths, from 3 mm to 5.5 mm, in steps of 0.5 mm. For each screw position, we used the CT to estimate the bone thickness (see below for more details), and used the next longer available screw length.

Importantly, even though the original tip of the screw was removed, we were still able to use these screws as self-tapping screws. However, it should be noted that the first few turns are more difficult than with un-modified screws, and special attention is required to make the screw find its way into the pre-drilled bone hole and to make sure the screw thread starts cutting into the bone. It might be helpful to practice this on a skull of a cadaver.

To measure the bone thickness for each screw location, the screw positions need to be defined in the CT reconstruction of the skull. For the milled version, one can directly overlay, in the planning software, the baseplate model with the skull reconstruction, and measure the bone thickness at each screw hole.

For the bent baseplate, different strategies can be used in order to infer the planned position of the screws. Following baseplate bending, one can mount the skull replica in the stereotaxic apparatus and measure the stereotaxic positions of the screw holes. The bone thickness can then be measured in the planning software at these pre-specified locations, after alignment of the skull reconstruction to the stereotaxic Frankfurt-zero. These measurements were done using one of the following software packages (see **Table 2**): 3D Slicer (Fedorov et al., 2012), Brain Voyager QX (Goebel et al., 2006), or ImageJ (Schneider et al., 2012).

One can also use the skull replica and the drawing of the baseplate outline to estimate the general location of the implant. Using landmarks on the skull, one can infer the respective CT slice and rough location of the screw. We have noticed that in most parts of the skull, the bone thickness changes smoothly, such that an approximate estimate of the screw position is sufficient.

Note that the measured bone thickness should then correspond to the length of the screw that extends below the baseplate leg into the bone; if at a particular screw position, the baseplate could not be perfectly adapted to fit the skull (primarily with the bent version), this distance needs to be added to the screw length.

### 2.3 A two-step implantation approach

#### 2.3.1 Headpost-baseplate implantation

For the baseplate implantation, following general anesthesia induction, the monkey is intubated and placed into the stereotaxic apparatus. The skin is shaved, thoroughly disinfected and the surgical site is surrounded with sterile drapes. Different approaches can be used in order to find the target location. For the milled baseplate, the stereotaxic coordinates are simply read from the CAD drawings. For the manually bent baseplate, a 3D printed skull replica can be fixed in a stereotaxic apparatus to read off the corresponding coordinates. Note that both approaches provide an initial positioning, yet the final adjustments (millimeter or less) are done manually in order to achieve the best fit to the underlying bone.

Another option is to use anatomical landmarks palpable through the skin for the approximate estimation of the position. For example, the frontal baseplate version is close to the supraorbital ridge. The distance from the supraorbital ridge to the intended position can be measured pre-surgically on a skull replica. In the surgery, the supraorbital ridge can be palpated, and the intended position thereby found.

Note that the baseplate is manually bent or directly milled to fit the skull, but the skull is covered with substantial muscle and skin, such that the baseplate seems to not fit until skin and muscle are removed.

Once the approximate baseplate position has been found, a sterile pen is used to draw a line on the skin to guide the skin cut. For frontal implants, a coronal incision is made that is placed about 1.5-2 cm posterior to the intended position of the central plate, to minimize the overlap between the later suture line and the implant. The skin is cut down to the fascia using a scalpel.

We use a periosteal elevator to detach the muscle from the skull (**Fig. 6A**), starting at its middle and frontal insertion and then working lateral and posterior. We minimize muscle detachment to the area covered by the baseplate legs and central plate. To this end, we repeatedly insert the baseplate underneath the partly detached muscle, onto the bone, to test whether muscle detachment is sufficient. Note that the periosteal elevator is only used to detach the muscle from the periosteum. In other surgeries, these instruments are typically used to clean the skull from the periosteum. However, in the baseplate implantation, we try to keep the periosteum intact, since it is the source of bone growth, respectively re-growth (Lin et al., 2014). It is crucial to keep the fascia and the skin moist during the whole procedure. They can be covered with gauzes that are regularly flushed with sterile saline.

**Figure 6.**
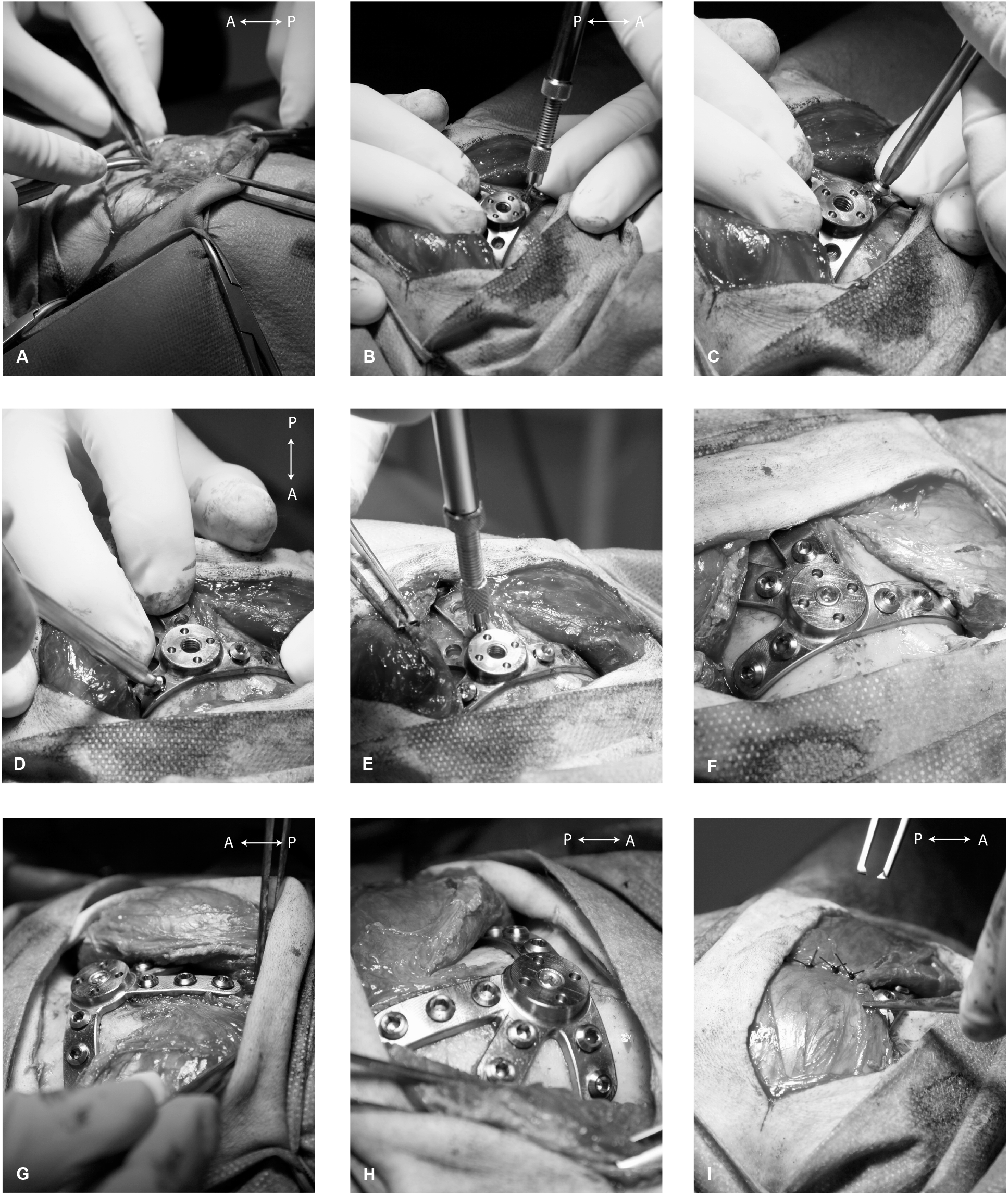
Implantation of headpost baseplate. **A**) The skin is cut and the muscle is detached laterally to expose the skull. **B**) When the implant position is found, the assistant firmly holds the baseplate onto the final location, and the surgeon drills the first bone hole using a manual drill combined with a drill-stop. **C**) The first bone screw is placed, and the same procedure is followed for each screw (**D-E**). Generally, the central screws are placed first, followed by the more lateral screws. Once two to three screws have been placed, the baseplate is sufficiently fixed to the bone so the assistant can release it. **F-H**) All bone screws are in place, and the cover screw has been added in the central plate. Note the essentially perfect fit of the CNC-milled implant to the skull. **I**) The muscle is brought back to completely cover the implant, and it is sutured together, followed by suturing of the skin. The approximate anterior (A) – posterior (P) orientation is indicated in the upper right corner of some panels. All photos in this figure show the implantation of Monkey H.

Once the skull is exposed, the baseplate position can be tested. Both the manually-bent and the milled baseplates fit so well to the skull that their final position is obvious simply from their fit. This is particularly evident for the milled baseplate. We experienced that even careful stereotaxic positioning was not able to place the baseplate into the position of optimal fit to the bone; therefore, after stereotaxic positioning, and drawing the pre-final position onto the bone, we removed the baseplate from the stereotax and left the final, minute, adjustment to the fit between implant and bone.

In case that parts of the underlying bone show sharp features (typically at the medial ridge), a small part of the bone can be slightly smoothed away by drilling. Unless really necessary, this step is avoided and the bone is kept as intact as possible. Note that these sharp features are typically visible in the CT. They can be taken into account during the planning procedure (only realistic for the milled version), or they need to be drilled away during the surgery.

In Monkey St, the medial ridge was very pronounced, which would have made it difficult to obtain a good fit of the bent baseplate. Therefore, during preparation of the bent baseplate, we smoothed the medial ridge of the skull replica using an electric drill (see **Table 4**). During the implantation surgery, we drilled the corresponding part of the bone ridge; to avoid removing too much of bone, this was done in several small steps, interleaved with fitting the implant to the skull, until the fit was optimal.

Once the optimal fit between baseplate and bone is found, the surgical assistant fixes the baseplate in this position by firmly pressing onto the central plate, and the surgeon adds the bone screws one by one. The bone screws are placed inside-out, i.e. starting with the screws close to the center (**Fig. 6B-E**) and then moving peripheral on the legs of the baseplate. Special care should be taken to insert the bone screws perpendicular to the skull surface to achieve the best possible grab and longer interaction surface. The bone holes for the screws are drilled with a manual hand drill combined with a drill stop (**Fig. 6B**) that prevents accidentally drilling too deep. The following procedure is followed:

1. We prepare the drill stop to accommodate the thickness of the baseplate. The length of the exposed drill is adjusted to correspond approximately to the thickness of the baseplate plus the expected skull thickness at the targeted screw position. In this, we take a conservative approach, aiming at leaving a very thin bone layer at the bottom of the drill hole.
2. The baseplate is positioned in its final position as explained above, and the drilling takes place through the holes of the baseplate. With the adjusted screw lengths, and the correspondingly adjusted drill depths, we found that we typically left the inner corticalis layer of the skull intact. This cautious approach has not led to any screw loosening (see Results, **section 3**).
3. We drive the screw into the bone and initially tighten it only loosely (**Fig. 6C-E**).
4. Once all screws are in place, they are fully tightened (**Fig. 6F**).

The screw hole in the central plate of the baseplate is then blocked with a cover screw (**Fig. 6F-H**) that prevents bone growth within it. One should make sure that this screw is sufficiently tightened against the bone, 1) to avoid that the screw is getting loose under the closed skin, 2) to ensure that the whole length of the screw hole is covered in order to prevent the bone from growing into this area.

When all bone screws are tightened and the cover screw is in place, the fascia and muscles are brought back above the baseplate and sutured together (**Fig. 6I**). In our experience, covering the implant with the fasciae and muscles improves the healing and prevents potential skin retraction following the later top-part implantation. The animals used in our experiments had quite extensive muscles, and this might have contributed to successful healing by forming a buffer between titanium and skin. Finally, the skin is sutured and the monkey stays on antibiotic treatment and painkillers for the following days. **Table 4** summarizes the implant parts and special instruments that are used in the baseplate implantation.

#### 2.3.2 Waiting time between surgeries

Following complete wound healing after the baseplate implantation, the top part is implanted. During this period, the baseplate implant is protected from the outside world, minimizing the danger of postsurgical infections that could jeopardize its osseointegration and the integrity of the underlying bone.

The waiting period between the two implantations differed substantially across monkeys (**Table 5**), ranging from 7.7 to 80.9 weeks (median: 37 weeks). The extent of the waiting period was mostly imposed by the needs and the progress of the respective experimental projects. Note that implant osseointegration takes 6-12 weeks (Hacking et al. (2012); also see discussion in **section 4.1**). Accordingly, we recommend a minimum waiting period of about 8 weeks, yet longer periods likely increase the probability and extent of osseointegration.

**Table 5.** Waiting time between headpost-baseplate and top-part implantations.

| Monkey | Waiting time (weeks) |
| --- | --- |
| Monkey L | 20.9 |
| Monkey Sk | 10.9 |
| Monkey D | 21.6 |
| Monkey G | 68.6 |
| Monkey Hu | 37.1 |
| Monkey Ch | 72.7 |
| Monkey T | 80.9 |
| Monkey St | 7.7 |
| Monkey M | 50.6 |
| Monkey H | 76.9 |
| Monkey C | 8.3 |
| Monkey K | 36.9 |

Importantly, during this period, there is no percutaneous implant and thus, no wound care is needed. We typically used this waiting period to train our animals in necessary procedures that do not require head-fixation, like chair training, acclimatization with the experimental set-up, initial head-free training in the recording booth.

#### 2.3.3 Top-part implantation

An important goal in this surgery is to produce a perfectly round hole in the skin that fits precisely around the top-part implant. In our experience, this is not feasible by manually cutting the skin, with or without adding sutures. Rather, we adopted an approach to punch a hole using a circular knife (“punch tool”; see **Fig. 7**). In this section, we present step-by-step the procedure and the tools we developed over the years. **Table 6** provides a summary of the implant parts and special instruments used in this implantation.

**Figure 7.**
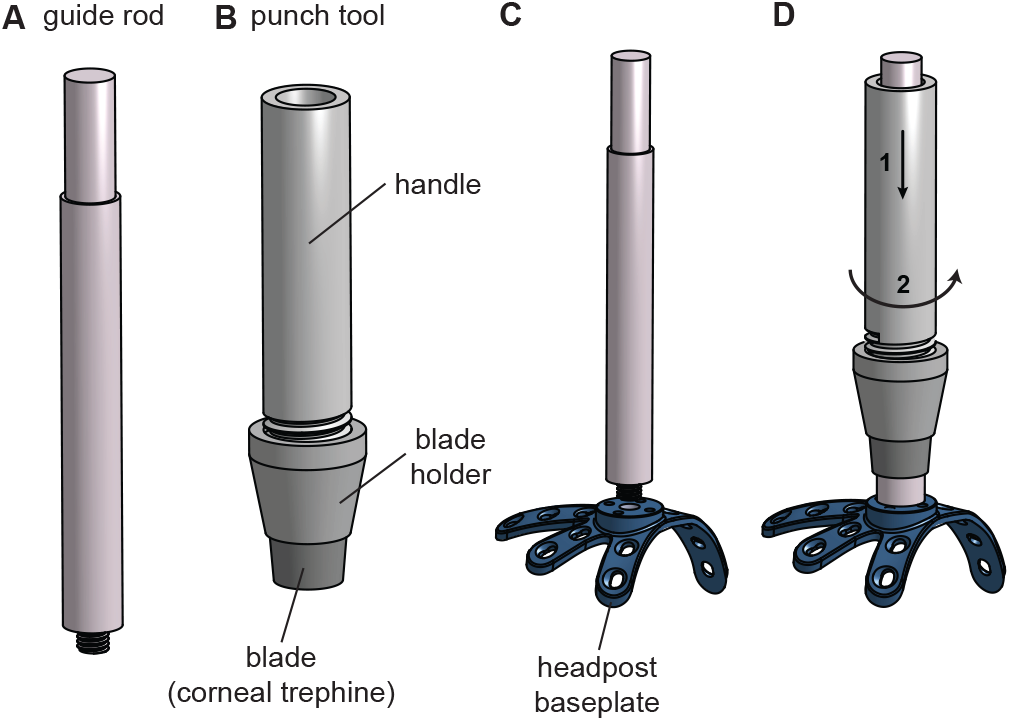
Illustration of the punch tool and its use in the top-part implantation. **A**) Guide rod. **B**) Punch tool. **C**) The guide rod is screwed into the central plate of the previously implanted baseplate. **D**) The punch tool slides on the rod to achieve a perfectly aligned and round cut through the scalp and muscle. Note that during the actual punching, the punch tool must not be merely pushed against the skin (illustrated with arrow 1), but it needs to be slowly rotated (illustrated with arrow 2). Different colors are used to illustrate the baseplate, the guide rod and the punch tool, even though they were all from metal.

**Table 6.**
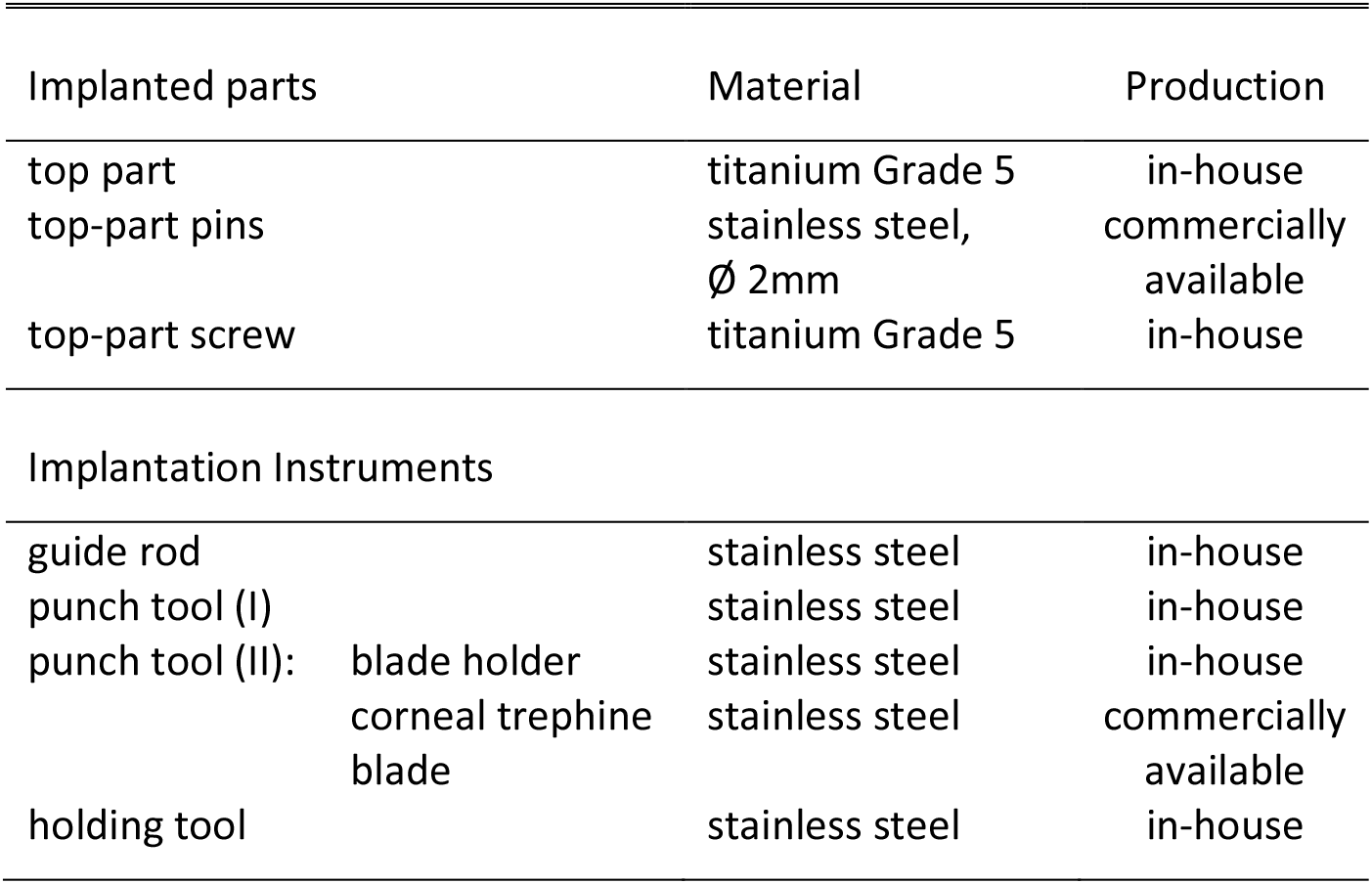
List of implant parts and implantation instruments used in the headpost top-part implantation.

The implantation of the headpost top part is a short surgery and can be performed under ketamine-medetomidine anesthesia and NSAIDs. Typically, the use of a stereotaxic frame is not necessary unless one needs to rely on stereotaxic coordinates in order to find the central plate of the previously implanted baseplate (different approaches are discussed below).

First, the skin is shaved and thoroughly disinfected and then, the central part of the baseplate has to be found. After shaving, one can typically see the healed suture line from the baseplate implantation which can help estimate the approximate position of the baseplate. If the baseplate is close to a bone landmark that can be palpated through the skin (like the supraorbital ridge), and/or if the muscle above the baseplate is thin, then the baseplate can be simply localized by palpation. If this is not the case, the central plate needs to be located by other means. In Monkey C, which was implanted with the central baseplate version, finding the central plate during the top-part implantation was difficult and time consuming, yet in the end successful. A way to facilitate this step could be to rely on stereotaxic coordinates; another option might be to leave a mark on the skin after the baseplate implantation, e.g. a tattoo; or position the central plate over the midline where typically there is no or only a thin layer of muscle. Placing the central part over the midline of Monkey K greatly facilitated this step. When the baseplate has been located, a small incision is made to expose the central cover screw, which is then removed (**Fig. 8A-C**).

**Figure 8.**
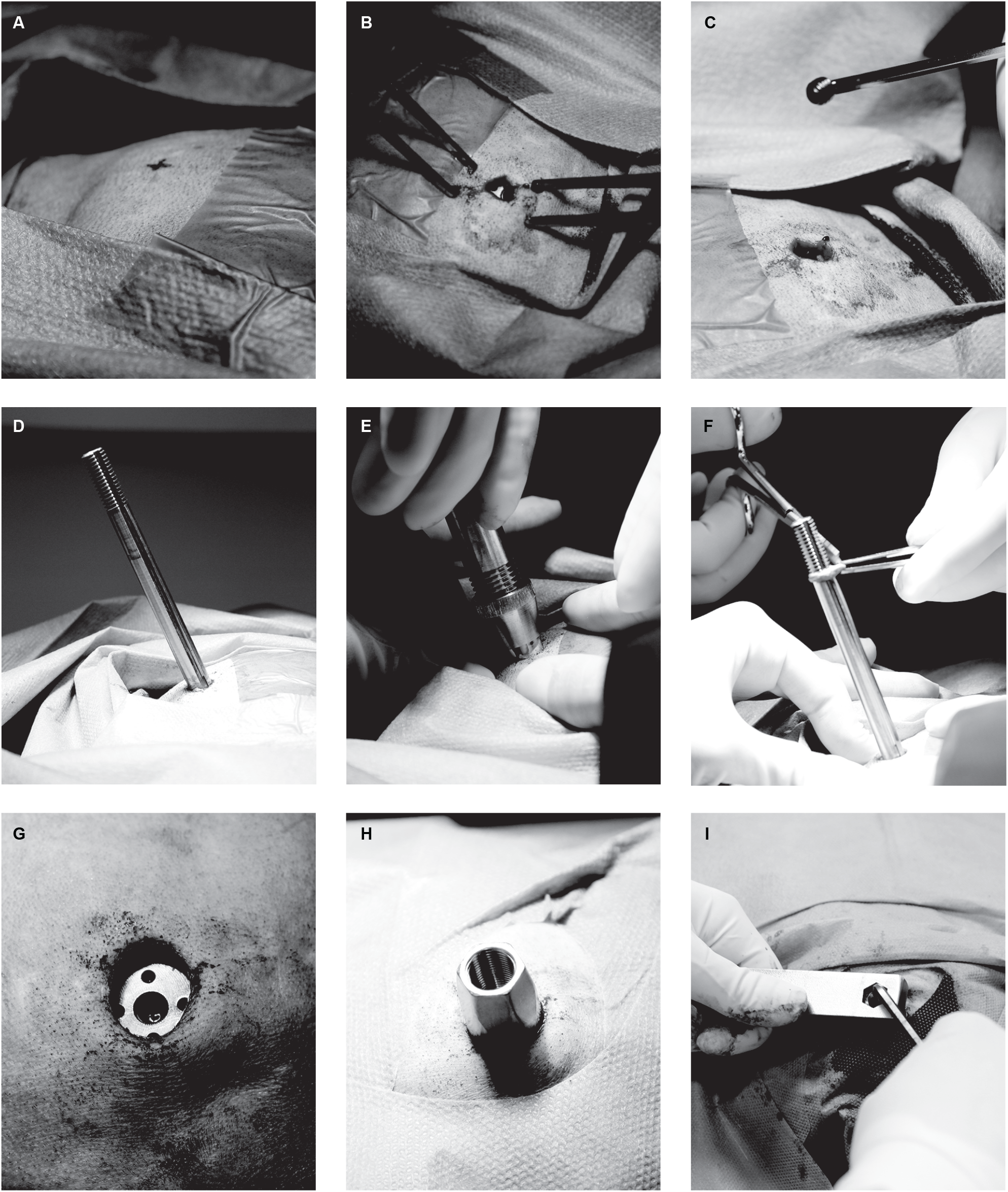
Implantation of the headpost top part. **A**) The central plate of the previously implanted baseplate is found, and a mark is made on the skin. **B**) At this location, a small incision through the skin is performed to access the cover screw. **C**) The cover screw is removed from the central plate, and **D**) the guide rod is screwed in this screw hole. **E**) The punch tool is sled onto the guide rod. While the assistant stretches the skin around the guide rod, the surgeon punches a hole into the skin by gently pushing and rotating the punch tool with the trephine blade. In the illustrated surgery, a corneal trephine blade was incorporated in a custom-made handle. **F**) The ring of cut skin is removed. **G**) The resulting hole in the skin is circular and centered on the central part of the baseplate. **H**) The top part is added. **I**) The central screw is tightened, while holding the top part with the holding tool. Panels A- H show the implantation of Monkey H, panel I shows Monkey C.

To perform the circular skin incision at the area where the top part will be implanted, we developed a custom-made punch tool with a circular sharp tip. For a similar result, a commercially available corneal trephine was integrated into the end of the punch tool (**Fig 7B** and **Fig. 8E**). A guide rod that fits within the punch tool is used to perfectly position the circular blade relative to the central part of the baseplate. To do so, the guide rod (**Fig. 7A**) is screwed into the central screw hole of the baseplate (**Fig. 7C** and **Fig. 8D**) and then, the punch tool can easily slide on it (**Fig. 7D**). Note that our version of this guide rod needs to be aligned very precisely to the baseplate to allow screwing it in. Finding the right angle might take a few minutes. This could be potentially improved by small modifications of the screw thread.

Before the punch tool touches down on the skin, the assistant stretches the skin in all directions away from the guide rod (**Fig. 8E)**. During the actual punching, the punch tool must not be merely pushed against the skin, but it needs to be slowly rotated, such that its circular blade can actually cut the skin and the underlying muscle, down to the central part of the baseplate. After the punching (**Fig. 8F**), the guide rod is removed (**Fig. 8G**).

Over time, we noticed some skin retraction around the top-part implant. To compensate for it, we use a punch tool that allows us to make a cut that is a bit smaller than the diameter of our implant, providing a tight fit. A trephine (Beaver-Visitec International, Waltham, Massachusetts, USA) with diameter of 11 mm was used in Monkey H for a top-part implant of 14 mm diameter. The use of the commercially available corneal trephine was inspired by a post in the NC3Rs Chronic Implants Wiki (NC3Rs, 2015) which presented their suitability for performing circular cuts through the scalp. Note that the corneal trephines are disposable and have to be discarded after a single use, because they are not sharp anymore. Similarly, our in-house punch tool becomes blunt during the process and needs to be resharpened after each surgery.

After performing the circular incision, the top part is implanted onto the baseplate (**Fig. 8H**), by inserting the two guide pins of the top part into the respective holes in the baseplate. The “top-part screw” is inserted and loosely tightened, such that there remains a little gap between the lower surface of the top part and the baseplate central plate. At this point, it is typically challenging to get all soft tissue (skin/fascia/muscle) out of this gap. We go around the top part with a pair of Dumont tweezers, at each angle pulling out any soft tissue. Subsequently, we apply a first, gentle, tightening of the top part.

For the final tightening, the head must not be in the stereotaxic apparatus. Otherwise, there is danger of damaging of the ears or the teeth. To be able to exert sufficient force for the final tightening, a holding tool with hexagonal cut-out is placed on top of the top part (**Fig. 8I**). A hex key is inserted into the top-part screw which is then tightened (by applying force between the holding tool and the hex key) with substantial force. Given the considerable force that is required, the tightening should not be done without the holding tool: in this case, the force would also be seen by the bone screws; by contrast, with the holding tool, the force is between the holding tool, the top part and the top-part screw, i.e. it remains within the metal structures of the implant. So far, we used a normal hex key and manual force estimation; yet the use of a torque screwdriver would likely be advantageous to ensure that the top-part screw is sufficiently tightened without applying unnecessary excessive force that could damage the thread.

### 2.4 Headpost holder

In order to increase handling safety, we developed and produced a headpost holder that allows remote posting (**Fig. 1C**). This holder is ≈31 cm long. The experimenter needs to touch and operate merely the proximal end, whereas the distal end attaches to the top part of the implant on the monkey’s head. In this way, the experimenter’s hands always remain at a safe distance from the animal’s head.

The holder, at its distal end, contains a central screw and a cap that fits onto the implanted headpost top part. When the headpost-holder screw is screwed into the top-part screw hole, this essentially pulls the hexagonal conus of the top part into the cap and thereby firmly secures the top part to the holder. **Figure 1C** graphically demonstrates an experimenter tightening the holder screw by turning the knob at the end of the holder. During this procedure, we found it useful to slightly wiggle the holder, such that the cap would “find” the optimal orientation to fit to the hexagonal conus of the top part.

The holder is produced by assembling the following independent pieces of stainless steel: 1) a “top-part cap” that fits onto the hexagonal surface of the headpost top part, 2) a tube, 3) a rod with a thread (M9 x 0.75) at its distal end, 4) a knob, and 5) a retainer cap. The top-part cap is welded onto the distal end of the tube, and the rod is inserted in the tube. At the proximal end, a retainer cap keeps the rod from slipping out while allowing it to freely rotate inside the tube. A knob welded at the proximal end of the rod allows the experimenter to comfortably rotate it during head (un)posting. The hexagonal shape of the top-part cap is cut using electrical discharge machining, because this achieves a perfect fit of the cap on the implanted top part with its fine and straight, i.e. non-rounded, edges.

In our setup, the headpost holder is secured to a headpost-holder mount (**Suppl. Fig. 3**), which is attached directly to the primate chair. This mount can be adjusted to hold oblique headpost-holder orientations, and thus, it accommodates a slightly oblique orientation of the headpost top part. This in turn allows to place the central plate of the baseplate away from the midline.

### 2.5 A three-piece connector chamber

#### 2.5.1 Design Considerations

Inspired by the two-piece headpost, we developed a connector chamber that consists of three separate pieces: a baseplate, a top part and a lid (**Fig. 9**). This modular connector chamber houses the connectors of multiple chronic electrode arrays and is implanted in a two-step implantation approach. The baseplate is the piece that comes into direct contact with the bone. It is implanted first and then covered with muscle and skin (see **section 2.5.3**). Following an adequate healing period of a few weeks that ensures initial osseointegration, the chronic microelectrode arrays are also implanted. During their implantation (see **section 2.5.4**), the baseplate is exposed and the top part that contains the array connectors is mounted and secured onto the baseplate. Finally, the lid is added to keep the connectors dry and protected outside of the experimental set-up.

**Figure 9.**
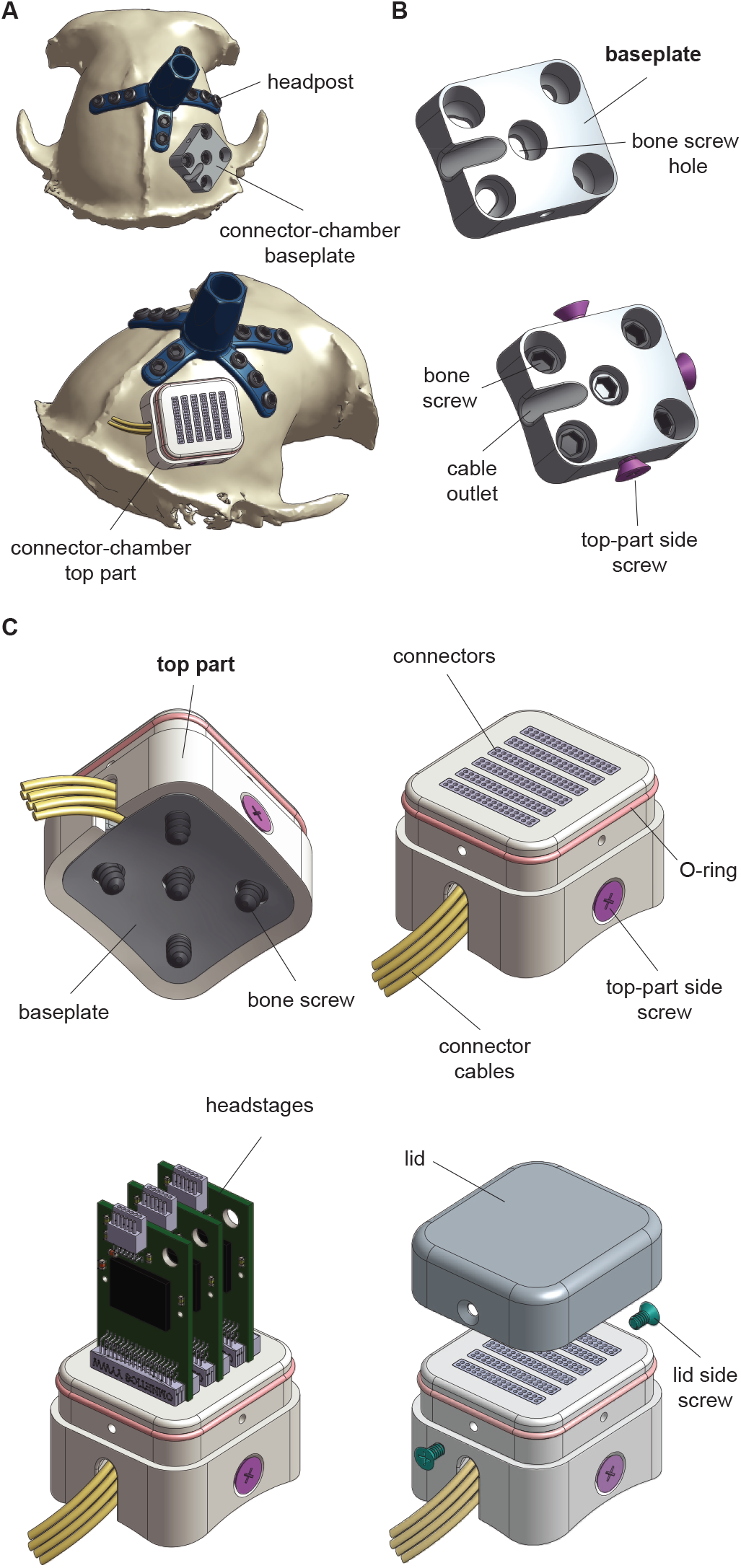
The modular connector chamber. **A**) Illustration of the two-stage implantation of the connector chamber (Monkey C). The baseplate is implanted first. Several weeks later, the top part that houses the electrode connectors is implanted in another surgery together with the electrode arrays (not shown here). Panels **B-C**) show in detail the baseplate and the overall implant, respectively. Note that all main parts of the implant (baseplate, top part and lid) are made of titanium; different colors are used for illustration purposes.

Two monkeys (Monkey C and Monkey K) were implanted with the central headpost-baseplate version. These animals are intended to be chronically implanted with microelectrode arrays in areas V1 and V4 of the left brain hemisphere. Each monkey would be implanted with six floating microelectrode arrays in total (Microprobes for Life Science, Gaithersburg, Maryland, USA), each consisting of 36 channels. The connector chamber was planned to be implanted over the right hemisphere, just posterior and close to the headpost baseplate (**Fig. 9A**) while it had to be large enough to house six Omnetics connectors (32-channel each; Omnetics Connector Corporation, Minneapolis, Minnesota, USA).

This type of arrays typically comes with a commercially available Crist Instrument titanium connector chamber (sometimes referred to as “pedestal”) that features radially extending legs that are fixed onto the skull with titanium bone screws. However, we had to develop a custom design for the following reasons; 1) these commercially available connector chambers could only house up to 4 Omnetics connectors, and 2) due to the proximity of the chamber to the headpost baseplate, the implant size had to be minimized. In fact, in none of the two monkeys there was enough space to fit a chamber with extending legs. Thus, we developed a new design that prioritized the reduction of implant size. This was achieved by planning an implant with the following main characteristics:

1. It is a footless implant. The titanium bone screws are incorporated within the connector-chamber baseplate (**Fig. 9B**). In other words, the size of the implant mainly depends on the number and the size of the electrode connectors that are planned to be housed within it.
2. It is a cement-free implant that is secured to the bone merely by five titanium bone-screws, without the need of an additional cement-cap. Such caps typically extend around the implant occupying considerable space on the skull.
3. Both the top part and its lid are mounted (to the baseplate and the top part, respectively) with side-screws (**Fig. 9B-C**), that do not significantly increase the implant’s footprint. In an earlier design version (that was not realized in the end), the top part was mounted on the baseplate with two vertical screws. By switching to side screws, the overall size of the baseplate was decreased from 39 x 23 mm to 21 x 19 mm. In addition, its thickness was reduced by 4 mm, which is beneficial for muscle and skin healing during the waiting period. This thickness reduction was possible, because the baseplate did not have to accommodate anymore the thread length of the vertical screws above the skull.
4. The implantation of the baseplate in a separate surgery significantly reduces the duration of the later electrode-implant surgery. Electrode implantations can be long and demanding, so they benefit strongly from deferring part of the efforts into a separate surgery.

#### 2.5.2 Planning and Manufacturing

The first planning step was the estimation of the minimum implant size that was required in order to fit the number of the connectors that were planned for our experimental needs. The connector chamber should house six 32-channel Omnetics connectors and allow enough space for three dual-64 channel headstages (Intan Technologies, Los Angeles, California, USA) to be comfortably connected onto them on a daily basis. For the top part, an additional wall thickness of 2 mm was included in our estimation. A rectangular box of these dimensions was fitted onto the skull model of the individual monkey and the three implant parts were subsequently designed: the baseplate, the top part and, the lid. All parts were produced from titanium Grade 2 and CNC-milled in a 5-axis machine. Here we provide a detailed description of each piece of the connector chamber.

Baseplate (**Fig. 9B**):

The bottom surface of the baseplate is planned to follow the geometry of the underlying bone, while its upper surface is flat in order to allow the top part to tightly sit against it. The baseplate incorporates five screw holes that are perpendicular to the bone surface and an outlet to allow the connector cables to leave the chamber. Finally, it includes three screw holes on the sides that allow the top part to be secured to it.

Top part (**Fig. 9C**):

The lower end of the top part slides over the baseplate, such that the baseplate almost completely disappears inside the lower end of the top part (a small gap between the lower surface of the top part and the bone was introduced to allow for potential bone growth). The top part is then fixed to the baseplate with lateral screws. Similar to the baseplate, the top part incorporates a cable outlet on its left side. On the upper surface of the implant, there are six slits that would allow the experimenter to connect the headstages to the electrode connectors.

The top part was first produced in-house and then, it was sent to the company that produced the electrode arrays (Microprobes for Life Science). They incorporated the Omnetics connectors into our top part and the free space was filled up with epoxy (**Suppl. Fig. 4A-B**). Subsequently, we applied a thin layer of bone cement (Super-Bond, Sun Medical Co. Ltd., Moriyama-shi, Japan) at the bottom of the top part, covering completely the epoxy (**Suppl. Fig. 4C** and **4E**), as a protection layer against fluids.

Three titanium side screws (M 2.5) fix the top part to the baseplate. During implantation, the cable outlet is also filled with Super-Bond to achieve proper sealing. An additional thin layer of cement (Super-Bond) is applied between the top part and the baseplate, preventing liquid from entering the implant.

The top part also features two pairs of screw holes for securing the lid. A spare pair of screw holes was added that could be used in case that the first set gets worn out after repetitive use.

Lid (**Fig. 9C**):

The lid is mounted on the top part with two titanium side screws (M 1.6). An O-ring is added at the interface between the top part and the lid to provide additional sealing.

#### 2.5.3 Implantation of the connector-chamber baseplate

The implantation of the connector-chamber baseplate is a very similar procedure to the headpost- baseplate implantation that has been extensively described in **section 2.3.1**. Here, we present the main steps of this surgery and indicate the main differences to the headpost implantation. Monkey C was implanted with the connector-chamber baseplate 28 weeks after the headpost baseplate and 19.7 weeks after the top-part implantations.

The location of the skin incision was defined based on the planned stereotaxic coordinates of the central bone screw of the implant. We tried to keep the size of the incision and the area of exposed skull minimal in order to make sure that the skin and the muscle surrounding the neighboring headpost would stay as intact and healthy as possible. The muscle was cut down to the level of the bone, and the skull was cleaned from connective tissue without removing the periosteum. The baseplate was inserted through the incision to test whether the size of the exposed skull was big enough.

The planned stereotaxic coordinates of the five bone screws were used to find the exact position of the implant. Note that this step differs from the implantation of the headpost baseplate, especially the milled version. In the latter case, one can use the stereotaxic coordinates as a first approximation to the target location. Then, thanks to its bigger size and long radially extending legs, one can manually adjust its position and find its optimal fit on the skull. In contrast, the connector-chamber baseplate is way smaller and footless which renders it infeasible to manually find its best fit. Instead, one needs to almost exclusively rely on the planned stereotaxic coordinates of the bone screws.

We used a sterile pen to draw the position of each of the bone screws based on these coordinates and then, we positioned the implant according to the marks and tested its fit against the skull. When the target location was found, the assistant held the baseplate in place and the surgeon started implanting the length-adjusted bone screws one by one, adding first the central screw and then the lateral ones. See **section 2.3.1** for a detailed description of the drilling and screw implantation steps.

Even though the bottom part of the connector-chamber baseplate was planned to follow the skull surface, its top surface was designed to be flat so that the top part can later on sit on it. This led to two important differences compared to the headpost baseplate implantation. First, when drilling using the drill stop, we had to find the proper angle that aligned the stop with the angle of the respective screw hole while ignoring the angle of the top surface of the implant. We found this step to be more difficult than in the implantation of the headpost baseplate. Second, the heads of the implanted screws were completely hidden within the screw holes. The gaps between the screw heads and the upper surface of the implant were filled with Super-Bond (see graphical illustration in **Suppl. Fig. 4E**) in order to avoid connective tissue growing there.

The top part of the connector chamber is mounted on the baseplate with three side screws. These screws were placed temporarily at the end of the baseplate implantation (as shown graphically in **Fig. 9B**), to avoid bone or connective tissue from blocking the screw holes. They have been removed and replaced by new ones during the top-part implantation (**section 2.5.4**). Finally, the muscle and the skin were closed up and sutured.

#### 2.5.4 Implantation of the connector-chamber top part and electrode arrays

During this surgery, the previously implanted connector-chamber baseplate is exposed, the chronic electrode arrays are implanted in the brain, and the connector-chamber top part housing the electrode-array connectors is mounted and secured onto the baseplate. Here, we provide a detailed description of the steps towards the implantation of the connector-chamber top part. For simplicity, we use in this section the terms ‘baseplate’ and ‘top part’ to refer to the main parts of the connector-chamber implant. The implantation is performed under general anesthesia. The monkey is intubated and positioned in the stereotaxic frame. Monkey C was implanted with electrode arrays and the top part 10 months after baseplate implantation (see **section 2.5.3**).

Following anesthesia induction, the surgical site was shaved and thoroughly disinfected. The previously implanted baseplate could be located by palpation. A large coronal incision through the scalp exposed both the left and the right hemispheres. This provided access to both the previously implanted baseplate (right hemisphere), and the area where the electrode arrays were to be implanted (left hemisphere). Part of the muscle covering the baseplate was cut and removed to expose the baseplate.

The three side screws (graphically depicted in **Fig. 9B**) on the baseplate, which were used to block bone and tissue growth within the screw holes, were removed, and the baseplate was cleaned with saline. There was pronounced bone growth around the baseplate. On some places, the new bone had almost reached the lower part of the side-screw holes. At this stage, the top part was mounted onto the baseplate in order to test its fit. Due to the extensive bone growth, the top part did not fit onto the baseplate and we had to remove some part of the new bone (using the Piezosurgery 3, Carasco, Italy). We repeatedly tested the fit of the top part onto the baseplate to limit bone removal to the required minimum. Note that even if the top part slides and fits onto the baseplate, it is still crucial to also test the fit of all three side screws that secure the two pieces together. We noticed later (see below) that even small misalignments due to the bone growth can prevent some of the screws to be properly mounted. The top part was then removed from the surgical area and stored in a sterile container.

Then, the muscle over the left hemisphere was detached from the skull using a periosteal elevator, the periost was removed, and a trepanation was performed. Then, the top part was placed onto the baseplate and the electrode arrays were implanted. The trepanation was covered with the bone flap, which was then sealed with bone cement (Super-Bond).

While mounting the top part onto the baseplate, we noticed that two out of the three side screws that were planned to secure the top part onto the baseplate (see **Fig. 9B**) did not fit, because we had only tested the fit of the top part, but we had not test-placed the screws. Some additional drilling allowed placement of the second screw. Further drilling was avoided to limit the time of dura exposure. To ensure proper stability of the top part, the empty space in the third screw hole was filled with bone cement. Bone cement was also applied to cover the other two side screws to ensure proper sealing. The cable outlet (see **Fig. 9B-C**) and all of the cables between where they exit the connector chamber and where they disappear under the bone flap were covered with bone cement. Finally, bone cement was also applied around the connector chamber to fill any small gaps between the bone and the top part.

The skin around the wound margin can often be swollen for some days or weeks postoperatively. In our case, this complicated the use of the connector-chamber implant during the first postoperative week due to: a) the overall low profile of the implant and, b) the use of side screws that secured the lid onto the baseplate. To solve this problem, some skin was removed one week later under a short ketamine-medetomidine anesthesia. This allowed us to securely open the lid in the next days. See **section 4.7.2** for a future refinement of our approach that will allow us to remove precisely as much skin as needed using a punch tool similar to that used in the case of the headpost top part.

### 2.6 Wound Care

We followed a similar wound care approach for both the headpost and the connector-chamber implants. Following the top-part implantation, the post-operative treatment was typically minimal. It included frequent hair-cuts around the wound margin and occasionally flushing with saline to clean the surrounding area. If the tissue looked irritated or slightly infected, the wound was flushed with antiseptic liquids. See **Figure 11** and **section 3** for later assessment of skin condition around the headpost.

## 3 Results

Twelve macaque monkeys were successfully implanted with headpost implants following our two-part design and two-step implantation approach. To date, there has been no implant failure or loosening from the bone in any one of the monkeys, in four cases more than 9 years after top-part implantation. The only failure that occurred so far was that in one animal (Monkey T), the top-part screw (located completely outside of the bone) partially loosened, such that the top part was not any more fully fixed to the baseplate, leading to small rotations. In this animal, we placed a new top part with new pins. This fixed the problem, and it did not reoccur. During the exchange, we found that the small movements of the top part had partially worn out at least one of the pin holes in the baseplate. Crucially, the baseplate remained firmly fixed to the bone. In order to prepare for any similar future cases, we then added two additional, spare, pin holes on the baseplate design. Yet, so far, we never had to use them. Note that in none of the animals, we observed any wearing out of the threads that connect the top part with the headpost holder, probably because these threads are relatively coarse.

**Table 7** summarizes the longevity of the headpost for all implanted monkeys. Implant longevity was defined as the number of years from the top-part implantation until today or until the end of the respective animal’s life (Monkey T was sacrificed, Monkey M died; both unrelated to the headpost). Monkey L has been successfully implanted for 9.7 years. In total, four out of the twelve monkeys have been implanted for more than nine years showing no implant-related problems. Monkeys H, C and K are the most recently implanted.

**Table 7.** Headpost implant longevity.

| Monkey | Years |
| --- | --- |
| Monkey L | 9.7 |
| Monkey Sk | 9.3 |
| Monkey D | 9.1 |
| Monkey G | 9.2 |
| Monkey Hu | 8.7 |
| Monkey Ch | 5.0 |
| Monkey T | 4.8<sup>***</sup> |
| Monkey St | 3.2 |
| Monkey M | 2.0<sup>***</sup> |
| Monkey H | 1.4 |
| Monkey C | 1.0 |
| Monkey K | 0.3 |

Number of years from top-part implantation until today or until sacrifice/death, the latter indicated with three stars (\*\*\*)

Nine monkeys underwent an additional CT scan following the baseplate implantation. These CT volumes were used to assess the long-term stability of the implanted bone screws. The time of assessment ranged from 0.1 to 7.9 years (**Table 8**). We found that at the time of the scan, there was no post-operative screw loss in anyone of the nine monkeys.

**Table 8.** Time of implant assessment

| Monkey | Years |
| --- | --- |
| Monkey L | 7.9 |
| Monkey Sk | 7.5 |
| Monkey G | 7.0 |
| Monkey Hu | 7.2 |
| Monkey Ch | 5.7 |
| Monkey T | 3.9 |
| Monkey St | 0.1 |
| Monkey M | 2.6 |
| Monkey H | 1.4 |

Years between the implantation of the headpost baseplate and the acquisition of the CT volume used for the implant assessment.

Monkey T was sacrificed due to reasons unrelated to the headpost. At that time, the baseplate had been implanted for 6.3 years. All bone screws were in place and high levels of osseointegration were observed above the legs of the implant (**Fig. 10**). Note that a bone screw was removed post mortem in the context of investigating the implant (see stars in **Fig. 10**).

**Figure 10.**
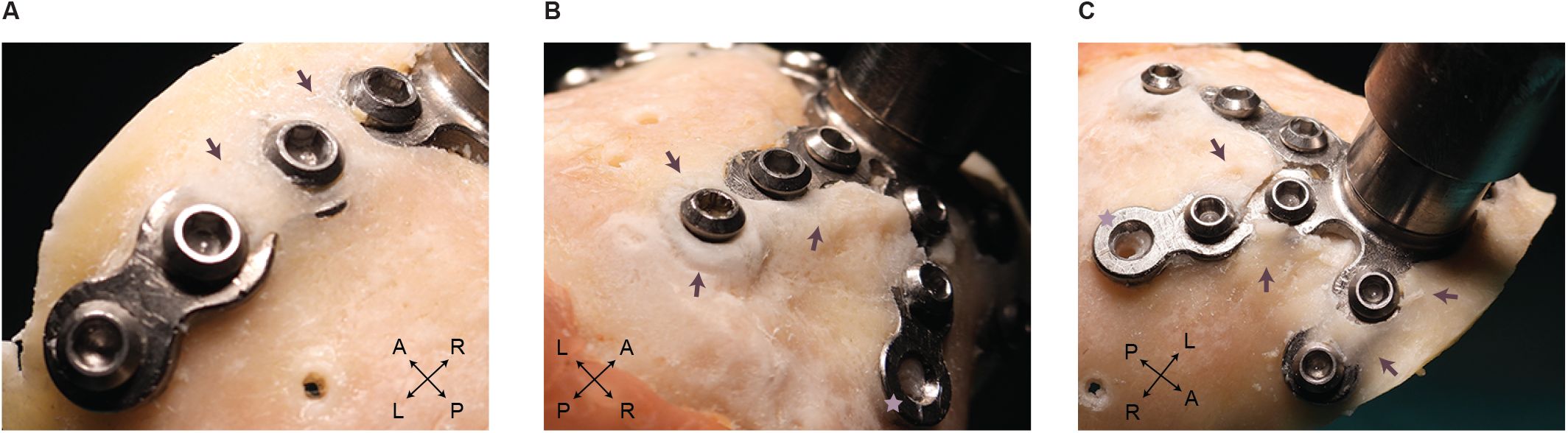
Osseointegration of the headpost baseplate. Pronounced bone growth over the implant legs 6.3 years after its implantation (Monkey T). Arrows point to areas of prominent bone growth. Stars indicate a bone screw that was removed post mortem. Bone holes outside the area of the implant were drilled post mortem for training purposes.

Different levels of skin retraction were observed across animals. **Figure 11** shows recent photos of the wound margin of three example monkeys. Skin retraction could not be assessed in monkeys M and T that were not alive at the time of evaluation. None of the ten monkeys showed extensive granulation tissue, or signs of infection like puss. Six monkeys showed no significant skin retraction, of which two example cases are shown in **Fig. 11A-B**. In Monkey G (**Fig. 11A**), the skin tightly followed the implant and looked dry and healthy even 8.7 years post implantation (when the photo was taken). Four of the monkeys implanted with the manually bent baseplate showed some level of skin retraction. Interestingly, in all of them the skin had almost exclusively retracted on top of the long anterior leg running over the left brain hemisphere. **Figure 11C** shows an example case of skin retraction (Monkey L), where the head of one bone screw has been exposed. Of the four monkeys with some skin retraction, three had only one bone screw exposed (Monkey D, Monkey Hu, and Monkey L), and one had two screws exposed (Monkey Sk). Importantly, in all four monkeys, the wound margin was dry, with no indication of ongoing retraction or bone exposure. The implants were stable and none of the animals showed any sign of discomfort.

**Figure 11.**
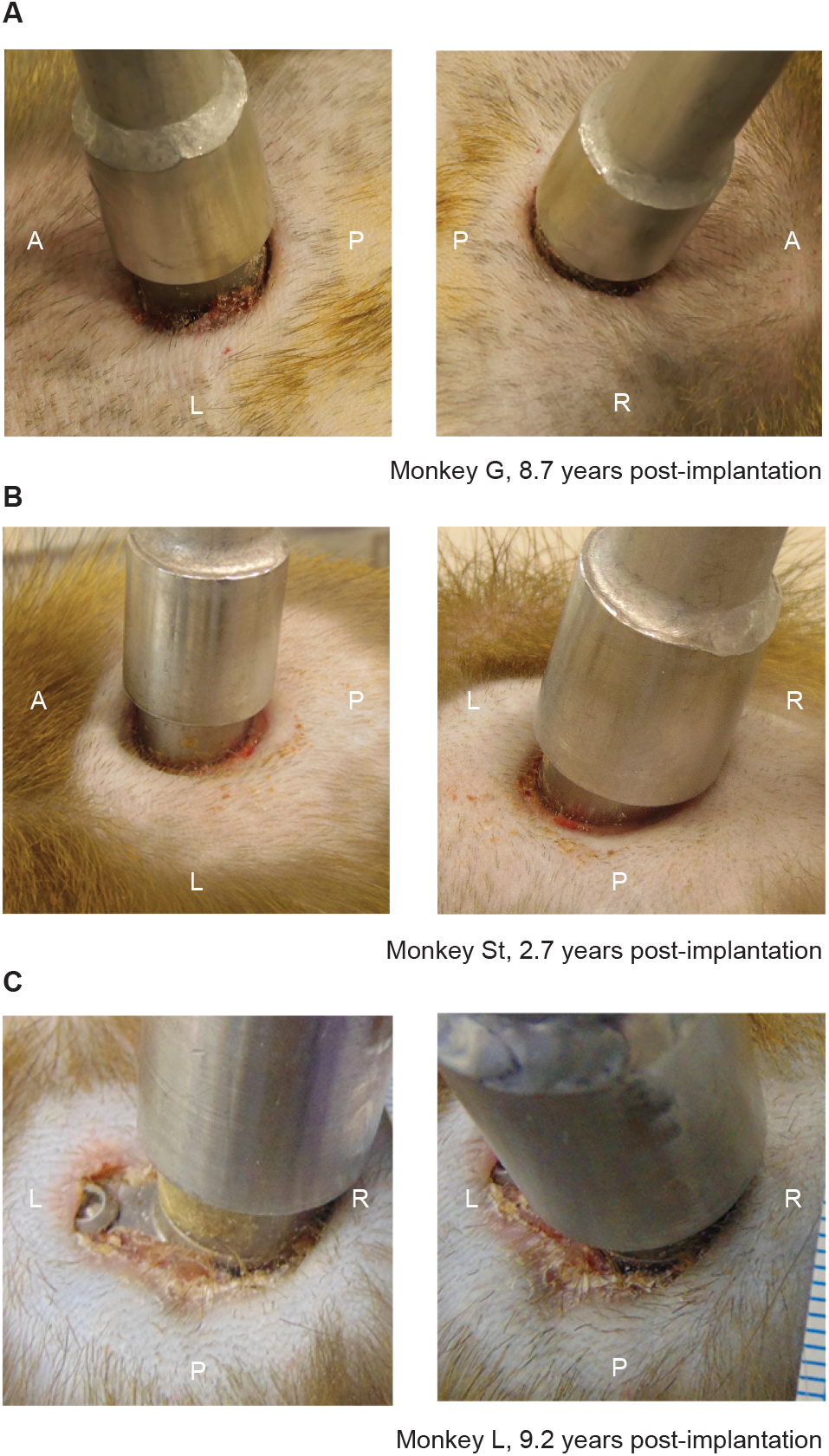
Skin condition around the headpost implant. The wound margins of three example monkeys are shown. **A-B**) Two example cases with no skin retraction. **C**) In Monkey L, the skin has retracted above one of the baseplate legs exposing the head of one bone screw. All photos were taken without prior wound cleaning on the same day. The anterior (A), posterior (P), left (L) and right (R) orientation is indicated with white letters on each photo.

While reviewing the post-operative CT volumes, we noticed that in two out of the four monkeys that showed some skin retraction, there was a small gap between the left anterior leg of the implant and the underlying skull (Monkey Sk and Monkey L; Note that there was no post-operative CT scan of Monkey D). This slight gap might have contributed to the observed skin retraction. However, skin retraction can be caused by many factors (see Discussion, **section 4.4**). Given that the implant longevity for these four monkeys with some skin retraction ranges from 8.7 to 9.7 years, we can conclude that the skin retraction has not affected the stability of the implant.

Three monkeys were implanted with headpost implants that were CNC-milled to closely match the skull surface. CNC-milling allowed us to improve the fit of our implants especially at the central plate of the baseplate. This central plate was difficult to fit to the skull in the version with the manually bent baseplate.

Finally, a novel connector-chamber implant was designed, produced and implanted in Monkey C. The baseplate was implanted first. Ten months later, the top part and the electrode arrays were implanted in a second surgery. During the top-part implantation, we observed: a) that the muscle covering the baseplate had completely healed since the baseplate implantation and, b) very pronounced bone growth around the baseplate. In several parts surrounding the baseplate, the new bone had grown for ≈2 mm reaching the side-screw holes. Parts of the new bone had to be drilled away to allow for the top part to slide and fit onto the baseplate. We expect that additional bone remodeling had taken place around the bone screws and underneath the baseplate which provides additional stability to the implant. At the time of writing this text (2.6 months post-implantation), the implant is stable, and the signal quality of the electrode arrays is good.

The skin around the implant is dry without signs of infection. The monkey shows no signs of discomfort. As expected, the skin has retracted over the bone cement that was applied to seal the cable outlet, exposing the bone cement in this area. There is no significant skin retraction around the rest of the implant. Due to the low overall profile of the connector chamber, we noticed that the lid and its side screws that secure it onto the top part might not be the optimal choice for the following reasons: a) the side screws are very close to the skin which makes it more difficult to remove them on an everyday basis and, b) because the lid is so close to the skin, it needs extensive cleaning every time it is removed. Based on this experience, we improved the connector-chamber design in order to surpass the difficulties we faced during implantation (see **section 2.5.4**) and to facilitate its everyday use. The new design is described in detail in **section 4.7.2**. Monkey K will be implanted with this new design.

## 4 Discussion

We described the planning, production and implantation of modular and cement-free cranial implants made of titanium. The modular nature of our implants allowed us to perform a two-step implantation approach that led to long-lasting, healthy implants even more than 9 years post implantation. To our knowledge, these are the longest follow-up cases reported for headpost implants. By reviewing post- operative CT scans, we also demonstrated the safety of using length-adjusted bone screws that are shortened to follow the thickness of the skull.

The two-step implantation approach combined with the introduction of a punch tool achieved a tight fit around the headpost top part without the need of additional sutures. This solved a frequent post- operative problem, namely the opening of sutures around the percutaneous part of the implant.

Overall, we describe several modifications with respect to previously developed methods, which likely contributed to our observations that the implants were surrounded by relatively irritation-free wound margins, and that they all lasted through the entire observation period. Together, those modifications therefore constitute an implementation of the 3R principles. In particular, we consider it a refinement if, in a given animal, the wound margin is free of irritation, and the headpost is long-term stable without the need of reimplantations. We also consider it a refinement that the screws are adjusted to the skull thickness, such that the risk of irritation or damage to the dura is minimized. Finally, we consider it a potential for reduction in animal numbers, if a given animal is in a healthy state with a stable headpost, such that it can potentially be used for further experiments.

We provide access to the drawings and models of our implants and the specialized tools that we developed over the years. We believe that sharing the methodological details and the long-term results from a significant number of animals can promote animal welfare by helping other labs to improve their methods.

### 4.1 The two-step implantation approach

Several previous studies have described a two-step implantation approach. For example, Pfingst et al. (1989) and Betelak et al. (2001) implanted titanium fixtures (anchor screws) that they let osseointegrate before securing cranial implants on them in another surgery. Blonde et al. (2018) implanted a large skull cap made of PEEK on which they subsequently added headpost and recording chamber implants. Finally, Chen et al. (2017) reported a two-step implantation of a footed titanium connector chamber.

Here, we further establish the suitability of this approach for the case of headpost as well as connector-chamber implants. The two-step approach allows to cover the implant with muscle and skin, protecting it from bacterial colonization. It also minimizes the risk of micro-movements of the implant relative to the bone, which might happen for one-piece implants with a percutaneous part, when the animal bumps with it against the chair or the home cage. Together, these factors likely promote the start of osseointegration, which seems essential for its later success. We think that the two-step approach was a crucial ingredient for the long-term stability that we observed for all implanted headposts, and we therefore recommend it as standard procedure. We estimate that the gain in animal welfare exceeds the “cost” incurred by the need for a second surgery; note that the top-part implantation is a short procedure that does not require intubation and typically lasts less than an hour.

Additionally, the two-step approach does not necessarily extend the overall time of preparation, because it is anyhow common practice to not place load on the implant for some period after implantation to allow sufficient osseointegration, also with one-step approaches. For example, Adams et al. (2007) waited two weeks, Hacking et al. (2012) six weeks, and Overton et al. (2017) four to twelve weeks between headpost implantation and first head fixation, depending on the age of the animal (see also **section 4.6**). Similarly, Lanz et al. (2013) mention that their monkeys were not headposted during the entire training period, until the recording chamber was implanted.

Adams et al. (2007) reported the formation of woven bone at the implanted area four weeks after the implantation of a footed, cement-free headpost implant. Hacking et al. (2012) showed new bone formation 14 weeks after the implantation of both textured and polished cement-free, titanium plates.

The two-piece design inherent to the two-step approach has a few additional advantages: The top part can be temporarily removed to fix other parts to the baseplate. We have used this option to place CT markers for stereotaxic calibration into well defined and reliable positions. The top part could in principle also be removed if an animal has a very long “holiday” or is not anymore used for experiments. Finally, the top part could also be exchanged for a modified piece if helpful for the experiment.

### 4.2 Shortened bone screws and the choice of screw type

The use of shortened bone screws did not compromise implant stability. In fact, we experienced no post-operative screw loss. Several other studies have supported the use of length-adjusted bone screws. Mulliken et al. (2015) presented a recording chamber in which the bone screws were counter- sunk into chamber walls such that only 2.5 mm of the screw thread entered into the skull. Similar to our approach, Pfingst et al. (1989) shortened bone screws according to skull thickness, and Overton et al. (2017) used commercially available bone screws that came in different lengths according to pre-surgical measurements of the bone thickness. In accordance with our experience, Betelak et al. (2001) reported that the skull thickness of macaque monkeys (*Macaca mulatta and Macaca nemestrina*) ranges from 2.5 to 4 mm, and they used a screw length of 3 mm (see also **section 4.6**). Note that we have successfully used the same screw planning approach (**section 2.2.5**) for the implantation of other implants like recording and chronic connector chambers, which we have so far embedded in bone cement (not described here).

The choice of the type of bone screw used can play a crucial role in the implant success. We believe that part of the long-term success of our implants was due to the use of the specific self-tapping bone screws employed here (**section 2.2.5**). We have experienced screw loss in earlier experiments (not described here) involving other types of screws. Note that screws can differ substantially with regard to the depth and shape of their screw thread.

### 4.3 Bent versus CNC-milled baseplate

We described two approaches to arrive at the individually shaped baseplate, namely the manual bending approach and the CNC-milling approach. Manually bent baseplates can be produced on 3-axis CNC machines from relatively thin titanium sheets in larger numbers, while discarding relatively little material, which makes them substantially cheaper. The CNC-milled baseplate version requires a 5-axis CNC machine and more planning and material, leading to substantially higher production costs. Together, these factors might limit availability of CNC-milled baseplates. Yet, while the manually-bent version provided an acceptable fit, this fit was essentially perfect with the CNC-milled version.

A promising alternative approach to CNC-milling is 3D metal printing. Titanium 3D printing has been successfully employed to produce headposts (Ahmed et al., 2022; Chen et al., 2017) and connector-chamber baseplates (Chen et al., 2017). The main advantage of 3D printing over CNC-milling is the lower production cost. However, note that 3D printing cannot so far produce the required precision for screw threads and very smooth surfaces. Therefore, in the case of our headpost implant, the following post processing would be required; 1) creation of internal screw threads at the central plate of the baseplate and at the top part, 2) smoothing of the external surface of the top part to avoid accumulation of dirt on its bottom part, and to ensure good fit to the headpost holder on its upper part. One potentially attractive option is to combine 3D printing of the monkey-individual baseplate (adding the central screw thread via post processing) with CNC-milling the top part, which anyhow remains constant. Yet, one noteworthy concern about 3D printed metal implants is that the metallic structure of the printed, i.e. successively deposited metal, is different from the structure of the metallic blocks that form the basis of milled pieces; how these structural differences affect suitability for use as implants will require further exploration.

### 4.4 Skin retraction

Similar to previous studies, the extent of skin retraction varied across monkeys (Overton et al., 2017), likely due to several factors: 1) The degree of manipulation of the wound margin by the individual monkeys, particularly during the days immediately following the top-part implantation; 2) Potential wound-margin infections at some point after top-part implantation; 3) The fit between the baseplate and the bone varied, with improvements in fit seemingly leading to reductions in skin retraction; 4) The degree of any early gap between the top part and wound margin, which we reduced by decreasing the diameter of the trephine below the diameter of the top part, leading to a tight fit both early after the top-part implantation and also later on. Regarding the latter point, note that in the NC3Rs Chronic Implants Wiki (NC3Rs, 2015), a similar approach is recommended, i.e. a 7.5 mm diameter trephine for a 10 mm diameter implant.

The combination of several refinements most likely contributed to the fact that our most recently implanted animals (Monkey Ch, Monkey St, Monkey H, Monkey C, Monkey K) so far show no significant skin retraction (range of years since top-part implantation; 0.3 to 5 years).

### 4.5 The three-piece, footless connector chamber

Chen et al. (2017) described a similar cement-free, two-step implantation approach of a footed connector chamber. The baseplate was implanted first and let to osseointegrate for several weeks. An important refinement in our design is the incorporation of the bone screws within the baseplate that led to a significant reduction of the overall footprint of the implant on the skull. Additionally, the footless design might reduce the risk of skin retraction that often occurs on top of implant feet (Mulliken et al., 2015).

Our connector chamber also features several improvements over the commercially available chambers that accompany the floating microelectrode arrays that we used. The commercially available options can only house up to four Omnetics connectors, are implanted in a one-step approach and are typically attached to the skull with bone cement. Our refined version can house six connectors. It is also friendlier to the bone thanks to its cement-free nature and the two-step implantation.

### 4.6 Considerations regarding the age and sex of animals

All animals reported in this study were male macaques that received headpost or connector-chamber implants during adulthood (range: 7 - 15.4 years of age). We estimate our approach to be similarly successful in the case of younger and/or female conspecifics. However, there are important differences between animals of different sexes and age groups that should be considered during implant planning.

For instance, in the case of young animals one should take into account developmental changes of skull morphology. Such changes are typically more pronounced in earlier developmental stages. To adapt the implant to the current skull shape, the preoperative imaging should happen close in time to the actual implantation. Longer waiting periods could lead to an imprecise fit. Overton et al. (2017) kept this time period to less than six months (range of ages at the time of implantation: 6.9-29.3 years). Interestingly, Chen et al. (2017) implanted partly juvenile monkeys (range at the time of implantation: 4-7 years) with a time period of up to ten months between implant planning and implantation. These authors did not report problems with the fit of the implants to the skull.

The skull thickness in the area of the implant is an additional factor that can affect the stability of implants and can vary with age and/or between sexes. Adams et al. (2007) compared the thickness of the frontal skull between a 26-month old male macaque monkey, two adult male and four adult female macaques. They report a mean skull thickness of 1.95 mm for the juvenile monkey, 2.93 mm for the male and 2.23 mm for the female monkeys. Based on these measurements, they concluded that one can safely implant juvenile monkeys with titanium headposts. They report the successful implantation of two juvenile monkeys (29 and 38 months olds) with one-piece headpost implants.

The adult male animals used in our experiments had quite extensive muscles (see **Fig. 6**), and this might have contributed to successful healing of the skin, by forming a buffer between the baseplate and the skin. Female and/or young animals might have less muscle. In any case, we recommend to cover as much area as possible of the baseplate implant with muscle and fascia. Even if parts of the implant are not completely covered by muscle, but only by skin, we expect the skin to heal normally, as long as the surgical site is free of infection.

At the same time, both female and/or younger animals typically weight less than their male and/or older conspecifics. Thus, the implant needs to sustain less overall load. In the case of young animals, as they gain weight with age, we expect that the osseointegration will also proceed and sustain more weight. Overton et al. (2017) waited longer periods before loading their headpost implants in the case of larger, stronger or geriatric animals and in cases that the bone might have been previously compromised. In our approach, one can accommodate these aspects by adjusting the waiting period between the implantation of the baseplate and the top part.

### 4.7 Future refinements

#### 4.7.1 Future refinements of the headpost implant

In the future, if the overall headpost position allows, we aim to place the central plate, and thereby the percutaneous top part, precisely on the midline. This had been technically difficult or impossible with the manually bent versions, yet has now become possible with the CNC-milled version and was the case in Monkey K (central baseplate version). We believe that this modification has the following advantages: 1) It avoids partly oblique top parts, requiring correspondingly oblique headpost holders, with corresponding technical challenges in fixing them to the chair; 2) It provides identical top part and thereby headpost-holder orientations across monkeys, easing the sharing of equipment across animals; 3) It allows to place the circular skin cut for the top part precisely into the midline, which is expected to be the ideal position with regard to the pattern of scalp vascularization, i.e. with blood supply running from lateral to the midline. While we plan to place the central plate on the midline, we will keep the legs with their screw holes away from the midline, to avoid screws from damaging the superior sagittal sinus. Note that in many animals, there is a bone ridge in the skull midline; if this ridge is very pronounced, it probably has to be drilled away at the position of the central plate.

Another potential refinement is the application of surface treatment on the implant baseplates to promote their osseointegration. Note that we did not apply any surface treatment to the titanium baseplates after CNC-milling and, for the manually bent version, after the manual bending. Thus, there was neither a polishing nor an extra roughening step. Surface roughening (Hacking et al., 2012) and/or coating with hydroxyapatite (Chen et al., 2017; Lanz et al., 2013; Ortiz-Rios et al., 2018) have been successfully employed before and might be considered to further promote osseointegration.

#### 4.7.2 Future refinements of the connector-chamber implant

Regarding the footless connector chamber, based on our experience from its first use in Monkey C, we have already implemented the following refinements for the implant of Monkey K:

In an attempt to minimize even further the risk for skin retraction, our future connector chamber is designed with a cylindrical shape, thus allowing for a circular skin incision to be performed, in a similar fashion to the method described for headposts in this paper, that is with a punch tool (see **section 2.3.3 and Fig.7**). We expect this improvement to lead to a better skin incision and to prevent the need for any corrective procedure (such as the one described in **section. 2.5.4**).

The baseplate side screws are designed to be inserted at a 25-degree angle above the skull surface, rather than horizontally. This will ease the handling of the screwdriver during surgery, allowing the screw to find its thread axis without having the screwdriver pressing too much downwards on the surrounding soft tissues.

As discussed in **section 3**, the cap will in the future be fastened exclusively from the top of the implant, without the use of any side screw. The O-ring will also be moved to the upper face, leaving the sides of the implant free of any design feature apart from the baseplate side-screw holes. We expect those smoother side walls to facilitate the daily cleaning procedure and improve the wound margin condition.

Finally, the cable outlet of the pedestal will be horizontal rather than vertical. The wires will therefore leave the pedestal side by side, as close as possible to the skull surface, minimizing the amount of bone cement needed to embed them.

These design changes have allowed to reduce the implant footprint to 531 mm^2^, to be compared with the 546 mm^2^ of the previous design iteration. The maximum baseplate height was reduced from 7.0 mm down to 5.6 mm.

## Supplementary materials

To facilitate the transfer of our methods to other laboratories, we share the models of the implants and tools that we developed (https://doi.org/10.5281/zenodo.7300042). **Suppl. Table 3** provides a summary of the shared files and formats. For an easy and interactive visualization of the models, open the respective “eDrawings” files (.html format).

## Acknowledgements

We thank the technical service team of the Ernst Strüngmann Institute and especially David Konietzny, Georg Haas, Thomas Bischoff, Joscha Schmiedt and Kai Rönnburg for their exceptional technical support and valuable contribution to all technical advances described in this work. We also thank Marianne Hartmann, Julia Hoffmann, Sabrina Wallrath, Olga Arne, and Johanna Klon-Lipok for their assistance in surgical procedures and animal care. We are grateful to Dr. Christa Tandi, Dr. Alf Theisen and Dr.Christiane Kiefert who provided excellent veterinary care and advice. We thank Alina Peter, Georgios Spyropoulos, Jarrod Dowdall and Tommaso Tosato who were involved in some of the experiments with the monkeys on whom this paper reports.

## Author contributions

**Eleni Psarou:** Conceptualization, Methodology, Investigation, Visualization, Writing- Original Draft, Writing- Review and Editing. **Julien Vezoli:** Methodology, Writing- Review and Editing. **Marieke Schölvinck:** Conceptualization, Methodology, Writing- Review and Editing, Supervision, Project administration, Resources, Funding acquisition. **Pierre-Antoine Ferracci:** Methodology, Visualization, Writing- Review and Editing. **Yufeng Zhang:** Visualization, Writing- Review and Editing. **Iris Grothe:** Methodology, Writing- Review and Editing. **Rasmus Roese:** Investigation, Writing- Review and Editing. **Pascal Fries:** Conceptualization, Methodology, Investigation, Writing- Original Draft, Writing- Review and Editing, Supervision, Project administration, Resources, Funding acquisition.

## Supplementary materials

### Supplementary tables

**Supplementary table 1.** Specifications of CT scans used in headpost planning per monkey.

| Brilliance Scanner |  |  |  |  |  |  |  |  |
| --- | --- | --- | --- | --- | --- | --- | --- | --- |
| Monkey Name | Scan Options | Protocol Name | Slice Thickness | Spacing between slices | KVP (kV) | X-ray Tube Current (mA) | Rows/ Columns | Pixel Spacing |
| M.L. | helix | Felsenbein kinder /Ear* | 0.8 | 0.4 | 90 | 53 | 1024 | 0.1953 |
| M. Sk. | " | " | 0.8 | 0.8 | 90 | 53 | 768 | 0.2357 |
| M. D. | " | " | 0.8 | 0.8 | 90 | 53 | 768 | 0.1745 |
| M. G. | " | " | 0.8 | 0.4 | 90 | 53 | 768 | 0.2422 |
| M. Hu. | " | " | 0.8 | 0.8 | 90 | 53 | 768 | 0.1927 |
| M. T. | " | " | 0.8 | 0.4 | 90 | 53 | 768 | 0.2031 |
| M. M. | " | " | 0.8 | 0.8 | 90 | 107 | 768 | 0.2604 |
| Planmeca Scanner |  |  |  |  |  |  |  |  |
| Monkey Name | Body Part Examined | Slice Thickness | Height | Width | KVP (kV) | X-ray Tube Current (mA) | Rows/ Columns | Pixel Spacing |
| M. Ch. | head | 0.4 | 500 | 500 | 90 | 5 | 500 | 0.4 |
| M. St. | " | 0.15 | 1068 | 1068 | 90 | 8 | 1068 | 0.15 |
| M. H. | " | 0.15 | 1068 | 1068 | 90 | 5 | 1068 | 0.15 |
| M. C. | " | 0.2 | 1001 | 1001 | 90 | 5 | 1001 | 0.2 |
| M. K. | " | 0.15 | 1068 | 1068 | 90 | 5 | 1068 | 0.15 |

\* The protocol has the German name "Felsenbein Kinder", which in English corresponds to "petrous part of the temporal bone in children"

**Supplementary table 2.**
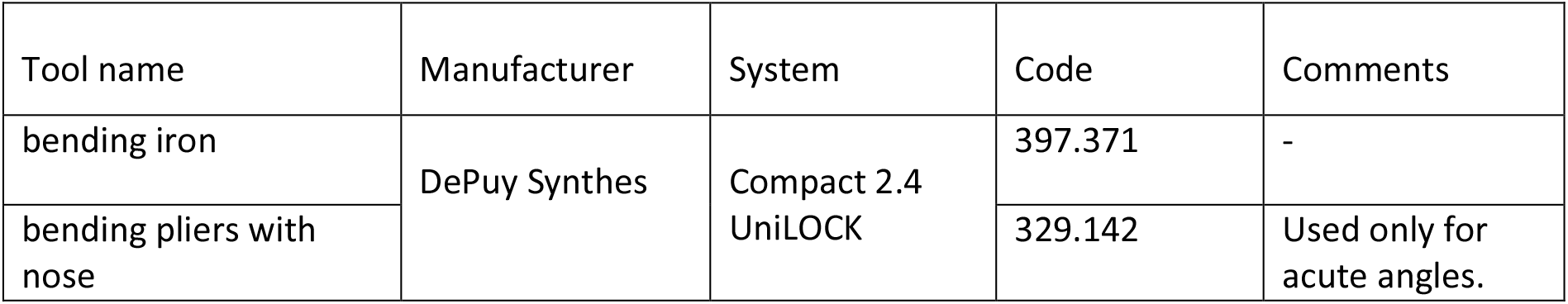
Bending tools used to pre-surgically customize the headpost baseplate.

**Supplementary table 3.**
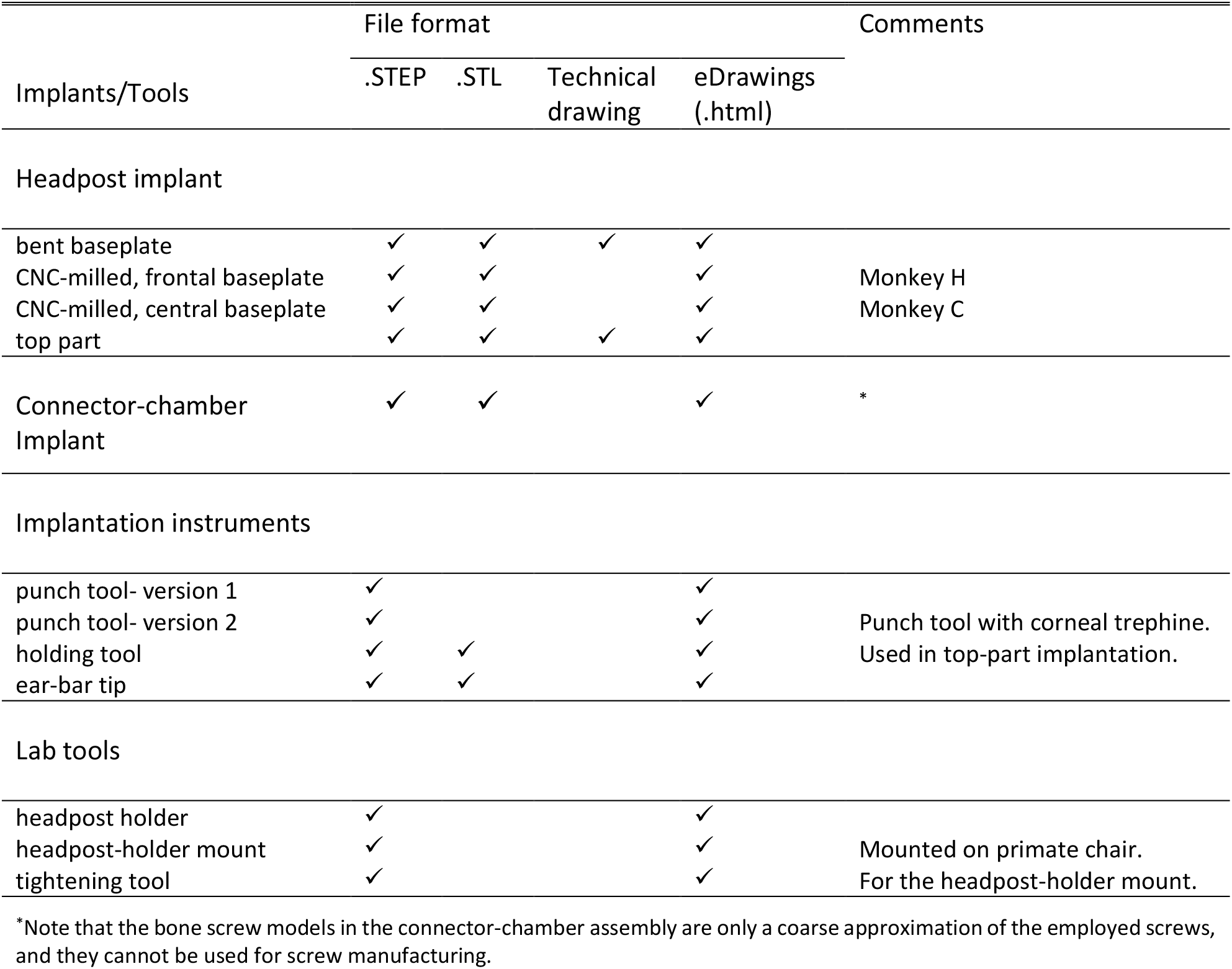
List of shared files.

**Supplementary Figure 1.**
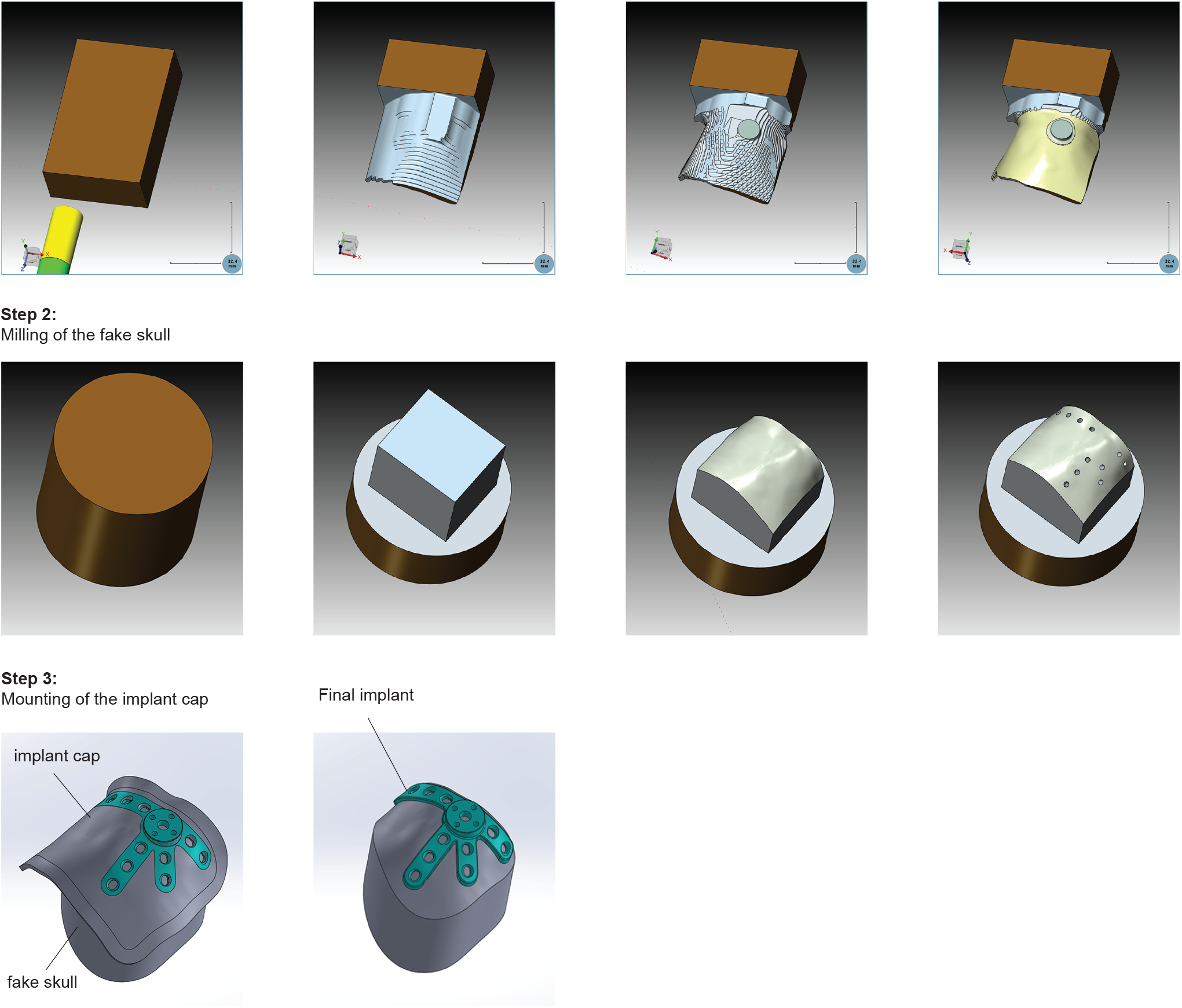
CNC-milling of a headpost baseplate using the “fake skull” approach. This figure presents a clamping approach that was used to CNC-mill the frontal headpost baseplate for Monkey H (see implant in Fig. 2B-C, Fig.3A and Fig. 6). **Step 1**: An “implant cap” was milled which incorporated the overall area of the baseplate implant. At this stage, the baseplate was milled in all details, except for its contour. Thus, the implant cap included: a) the precise geometry of the bottom and upper implant surfaces, b) the central plate and c) the bone screw holes (as shown in Step 3). **Step 2**: In order to ensure proper clamping and support of the thin implant cap while milling its implant contour, we devised the following approach: A head model (referred to as “fake skull”) was produced 1) which served as a base on which the implant cap was mounted, 2) which provided sufficient support to the implant legs while milling their contour, 3) which was easily clamped. The fake skull was milled from stainless steel and featured screw holes that were used to attach the implant cap on it using its bone-screw holes. **Step 3**: The implant cap was secured on the fake skull and its contour was milled. The final baseplate implant was then released from the fake skull. With this approach we were able to produce a headpost baseplate that showed an excellent fit both on the 3D skull replica of the monkey (Fig. 2B-C) and on the real monkey skull during implantation (Fig. 6). Nevertheless, it had two main disadvantages; 1) it was time consuming because it required the planning and milling of an additional piece (fake skull); 2) special attention had to be given to avoid imprecisions that could emerge while re-clamping the implant cap using the fake skull. For these reasons, we moved to the simplified approach presented in Supplementary Figure 2.

**Supplementary Figure 2.**
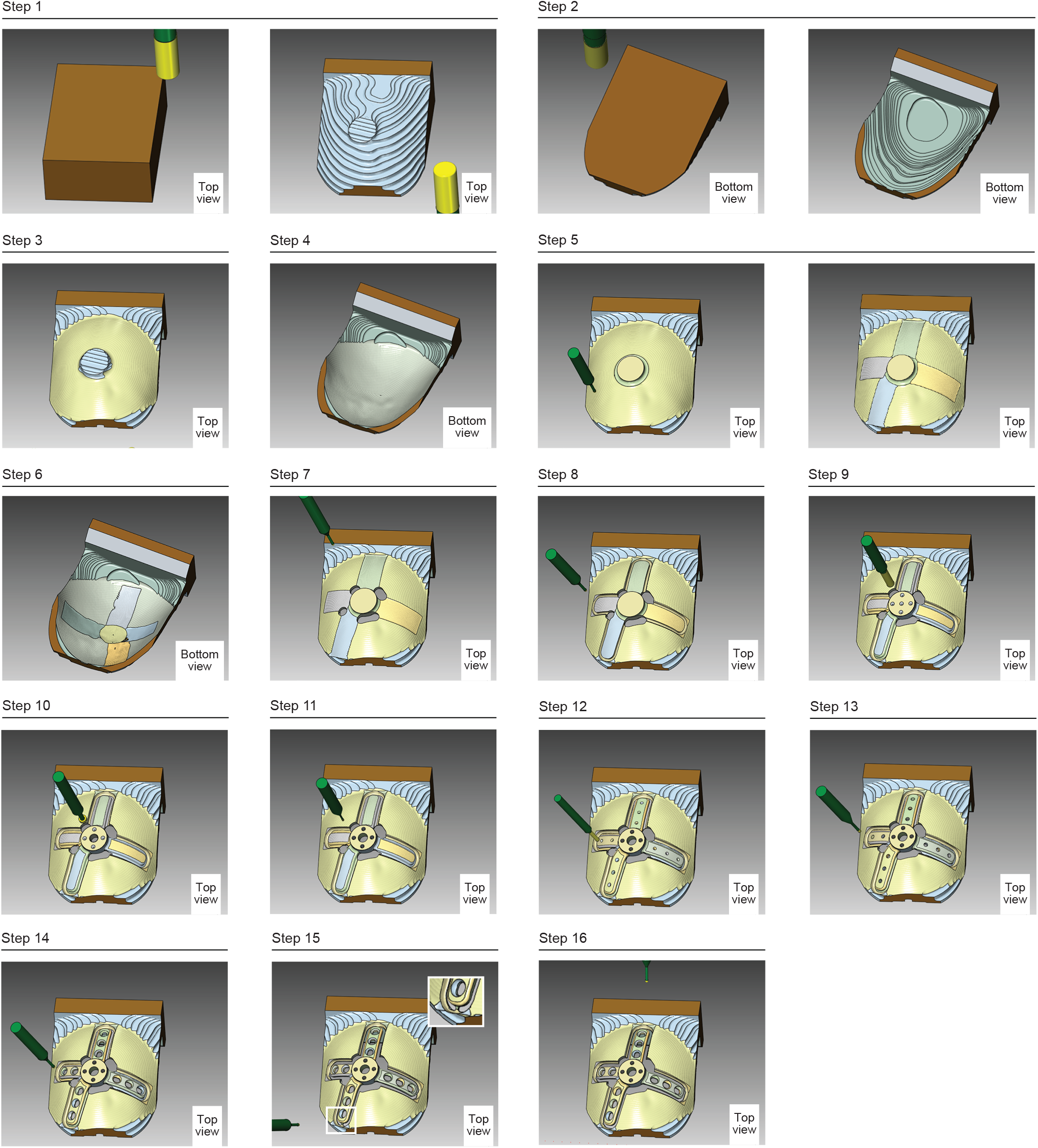
CNC-milling of a headpost baseplate without a “fake skull”. Graphical illustration of the CNC-milling steps of a headpost baseplate without the use of a “fake skull”, illustrated here for the central headpost-baseplate version (see Fig. 3B). The CNC-milling procedure contained the following steps: (**1-2**) A block of titanium was clamped from one side and a relatively thick cap was milled on the top and then on its bottom surface; this cap coarsely followed the outer skull surface in the overall implant area. More precisely, the cap corresponded to this surface plus 0.5 mm of material on the concave surface, and plus 0.5 mm of material on the convex surface. (**3-4**) Additional milling produced a more detailed surface (target implant surface plus 0.1 mm on the top and bottom surface). (**5-6**) In the next steps, the area of the implant was precisely milled to match the final surface. (**7**) The contour of the implant central plate was milled and then, (**8**) the upper part of the contour of the implant legs. (**9-11**) The central screw hole and the pin holes of the central plate were drilled. (**12-14**) The bone-screw holes were milled. (**15**) The baseplate was then partially detached from the cap by milling along its contours, while leaving four bridges, one bridge between each baseplate leg and the cap (see inset). These bridges ensured that the implant stayed attached to the cap. (**16**) In a final step, those bridges were milled away; to keep the baseplate in place during this procedure, high-performance tape (gaffer tape) was applied away from the bridges.

**Supplementary Figure 3.**
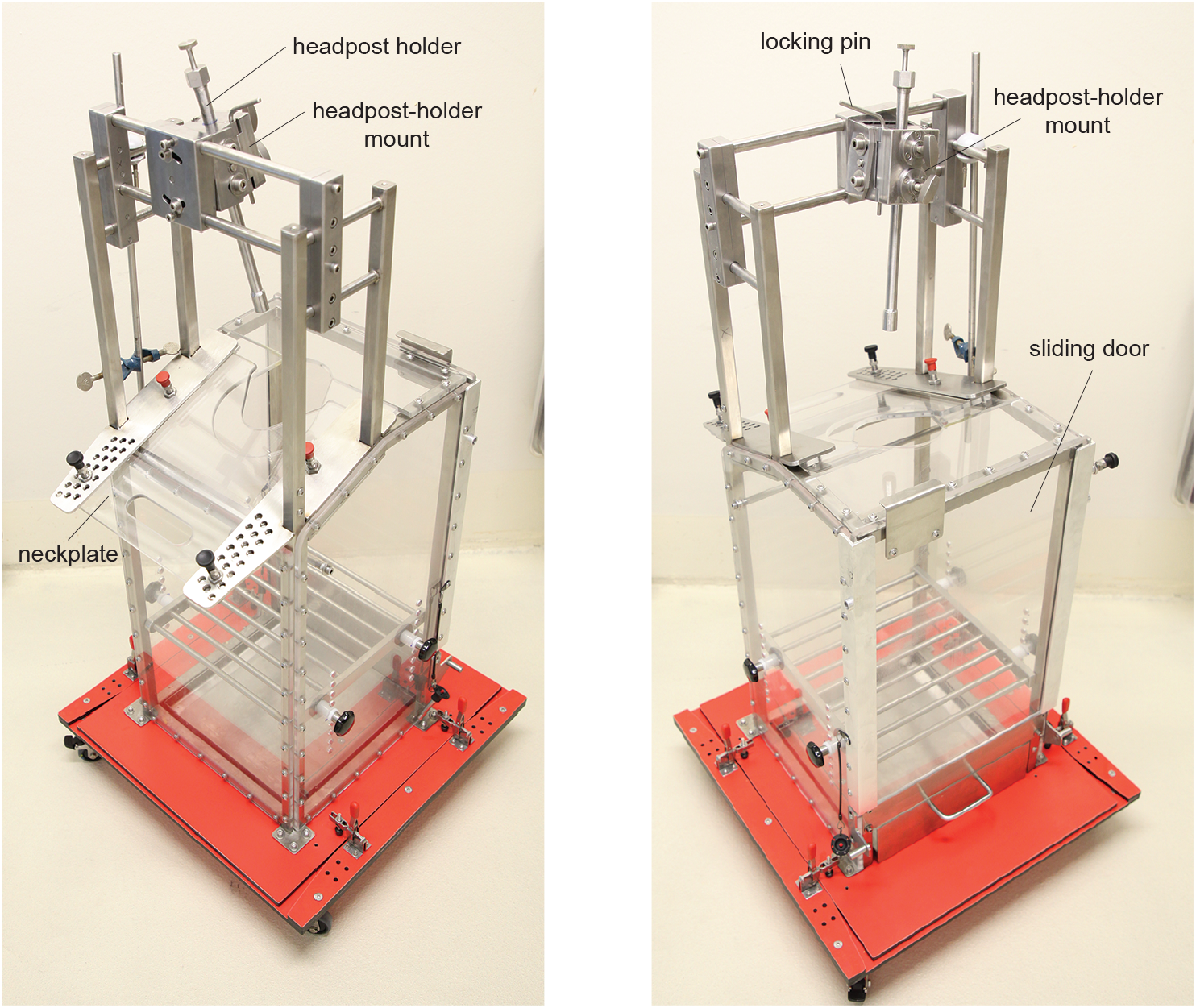
Primate chair and headpost-holder mount. Photos of the primate chair and the headpost-holder mount. The headpost holder is secured to a mount that is attached to the primate chair. The orientation of the mount can be adjusted to accommodate oblique headpost-holder orientations.

**Supplementary Figure 4.**
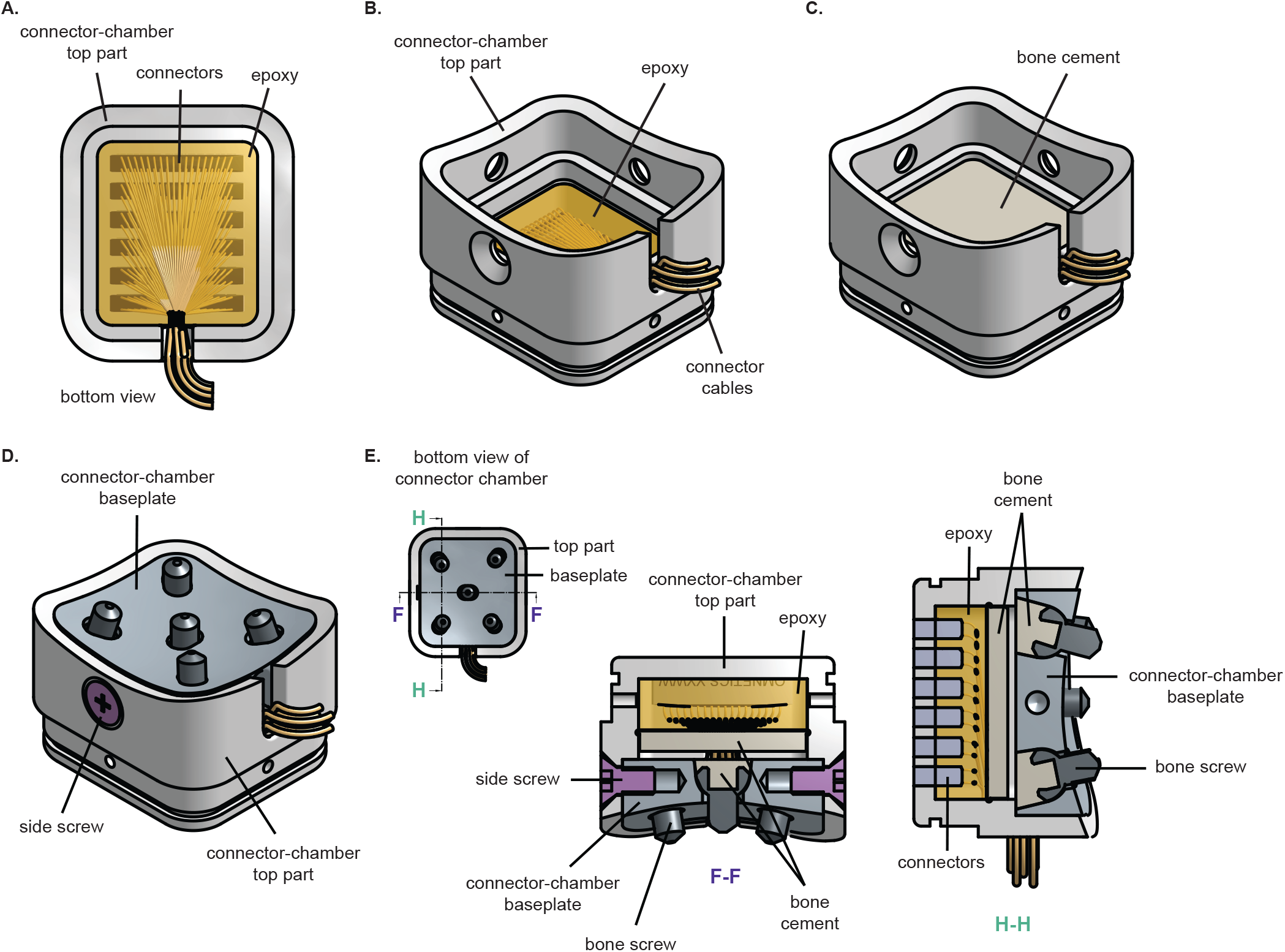
Sealing of electrode connectors in the connector-chamber top part. **A-B**) The Omnetics connectors were incorporated in the top part, and the free space was filled with epoxy. **C**) Subsequently, the epoxy was covered with a thin layer of bone cement to provide an extra layer of sealing against fluids. **D**) The connector-chamber top part is shown mounted onto the baseplate. **E**) Cross-sections (F-F and H-H) of the connector-chamber implant.

